# Revealing imatinib-kinase specificity via analyzing changes in protein dynamics and computing molecular binding affinity

**DOI:** 10.64898/2026.02.02.703340

**Authors:** William Troxel, Emily Vig, Chia-en Chang

**Author notes:** Correspondences Chia-en Chang.

## Abstract

Drug promiscuity is a double-edged sword where a small molecule acts on multiple biological targets to induce toxicological or therapeutic benefits. It is possible to exploit promiscuity to expand treatment options without the prohibitive costs of designing a new drug. Imatinib is a representative case, exhibiting varied affinities and inhibitions to different kinases. It binds most favorably to Abl and Kit kinases, intermediately to Chk1 and Lck kinases, and least favorably to p38α and Src kinases. The strongly conserved features of the ATP-binding site render imatinib’s molecular binding determinants unclear despite over 25 years of interrogation. To address this question, molecular thermodynamics, force distribution analysis, residue sidechain dihedral correlations, and principal component analysis were computed using trajectories from all-atom molecular dynamics simulations in explicit solvent. The results of these simulations agree with experimental affinity and binding data, enabling highly predictive factors for imatinib’s binding specificity from free- and bound-state simulations through a global protein network of protein-ligand interactions, changes in sidechain dihedral correlations, and shifts in the secondary motifs modulating binding site access corresponding with well-characterized kinase “breathing motions.” The sidechain dihedral correlation network also identifies distal mutants known to reduce patients’ imatinib sensitivity. Higher imatinib-kinase affinity trends with a loss in sidechain dihedral correlations and diminished secondary motif migration following binding, corresponding with more restricted configurations, to reduce solvent approach and ATP competition. Lower-affinity proteins show enhanced sidechain dihedral correlation and exaggerated secondary motif motions. This is consistent with a tendency to expose the protein pocket, facilitate solvent entrance, and increase ATP competition. Using imatinib as a model system, this study shows residue correlation, force interaction, and essential principal components can effectively forecast imatinib-kinase binding specificity and introduces an effective approach to repurpose and design high-affinity binders for off-target applications more generally.

## Introduction

Drug promiscuity, the ability of a single compound to interact with multiple biological targets, has emerged as an important phenomenon in drug discovery due to the resulting medicinal and toxic side effects.^1,2^ Promiscuity is controlled by molecular recognition, the specific non-covalent interactions that form between two or more molecules when they approach each other, such as when a small molecule inhibitor binds to a protein kinase.^3^ Rational drug design (RDD) leverages molecular recognition by creating small molecules that target, bind to, and inhibit specific proteins to achieve therapeutic effects.^3–5^ Issues emerge when multiple proteins share similarities in their binding pockets, such as protein kinases, where one drug targets the conserved ATP-binding site and binds to multiple kinases despite a wide variance in their primary sequences, introducing unexpected side effects.^3,6–8^ Simultaneously, if one compound binds to multiple different proteins, it may improve its disease-treatment potential while bypassing the expensive process of designing new drugs.^9^ While traditionally viewed as an undesirable source of off-target toxicity, promiscuous binding can explain the success of repurposed drugs. Understanding the molecular and structural factors that govern drug promiscuity is therefore critical for both predicting adverse interactions as well as guiding RDD.

Streamlining the drug prediction process by using novel analysis methods on drug-bound kinase data has great potential for basic research, translational applications, and commercial success. Tyrosine kinases (TKs) and serine/threonine kinases (STKs) modulate cell signaling pathways, making them attractive therapeutic targets for anti-cancer drugs.^9^ They are the second most targeted druggable macromolecule, only after GPCRs, with 25% to 33% of international drug development programs targeting kinases.^10,11^ Over 60 known kinases are targeted by FDA-approved drugs, and according to the NIH Illuminating the Druggable Genome project, there are more than 160 understudied targets.^12–15^ Highly targeted binding necessitates RDD to craft inhibitors to affect one biological target of interest, such as imatinib with Abl kinase, amidst high binding site similarity shared with multiple other kinases (Figure 1A). Imatinib was the first TK inhibitor designed to target one specific protein, the Bcr-Abl gene fusion protein. Nevertheless, since its approval in 2001, it has been discovered to target multiple different kinases despite primary sequence variance across proteins (Figure 1B). It is a 2-phenylaminopyrimidine derivative with pharmacophores to enhance oral bioavailability, TK binding, and Bcr-Abl selectivity (Figure 1C).^16^ These off-targets include Kit kinase to treat gastrointestinal stromal tumors, platelet-derived growth factor receptor to treat dermatofibrosarcoma protuberans, and colony-stimulating factor 1 receptor to mitigate rheumatoid arthritis-related bone damage.^8,16–18^ Recent proteomics studies reveal significant changes in ATP-binding capacities for Chk1 kinase in K-562 cells, involved in the DNA damage response, suggesting it may be a novel target for imatinib treatment.^19^ These pharmacological uses show that numerous unanswered questions remain about molecular determinants influencing imatinib’s binding specificity to protein kinases, and how pertinent it is to uncover these factors to expand targeted applications.

**Figure 1.**
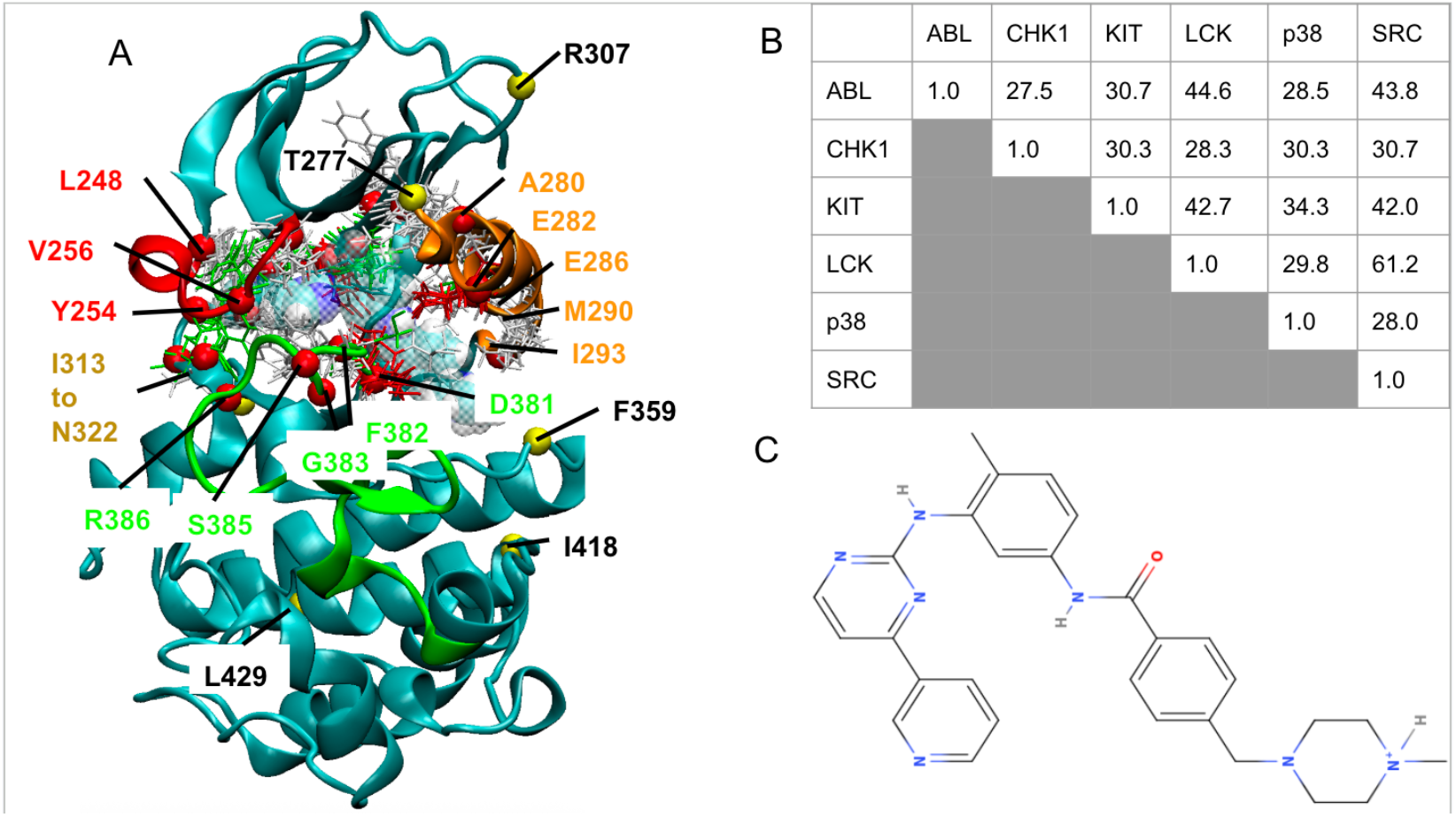
**A:** Superimposing the kinase domains of Abl (Cyan NewCartoon), Chk1, Kit, Lck, p38, and Src. Imatinib is shown with Van der Waals representation in the trans configuration. Binding site residues are marked with the red spheres. Distal residues from the binding site found as part of a key correlation network are marked with yellow spheres. The phosphorylation loop (P-loop) is shown in red. The alpha C-helix (αC-helix) is shown in orange. The activation loop (A-loop) is shown in green. These three motifs constitute the catalytic triad. The colored licorice residues show the high homology of the binding site residue sidechains identified using GROMACS force distribution analysis. **1B**: Pairwise percentage of sequence identity of Abl, Chk1, Kit, Lck, p38, and Src based on multiple sequence alignment from the NCBI blastx program. **1C**: 2D protonated imatinib schematic drawn with MolView.

Three decades after imatinib entered the market, a satisfactory answer still has not been obtained to explain its differential affinities for specific kinases. Early crystallography showed that the Abl and Src kinases, with a 3000-fold affinity difference, could adopt similar inactive configurations to each other, indicating imatinib selectivity is not necessarily defined by the static inactive conformation, and that protein kinase dynamics contribute.^20,21^ A crystallographic investigation of Abl, Kit, Lck, and p38α kinase domain changes due to imatinib binding found an apparent increase in solvent exposure correlated with lower imatinib-kinase affinity.^22^ Different hypotheses emerge for either small configurational changes influencing imatinib binding, probabilistic differences of Abl and Src adopting the inactive configuration, or induced-fit configurational changes after binding, demonstrating the challenges of elucidating the imatinib-kinase specificity question.^20,23^ Since crystal structures do not show system dynamics, computational methods are necessary to bridge the gap and provide insights into how these kinase systems move over time.

Molecular modeling provides unique insights, including quantifying drug-protein binding energetics to enable comparisons between several distinct kinases binding a drug, and interrogating differences in drug binding/unbinding with wild-type and mutant proteins. Computational methods, such as free energy perturbation (FEP), were used to interrogate the absolute binding free energies of imatinib with Abl, Kit, Lck, and Src kinases, elucidating key noncovalent interactions that compensate for the loss of imatinib’s degrees of freedom following binding.^24–27^ Recent studies have investigated microsecond to millisecond timescale simulations to examine specificity trends for wild-type and mutant Abl.^28–30^ However, the selectivity of imatinib toward multiple proteins remains insufficiently characterized in the current studies. The challenges surrounding computational models indicate that further investigation and alternative approaches are necessary for a comprehensive understanding of imatinib-kinase specificity.

This study utilizes all-atom classical molecular dynamics (MD) simulations with various post-analysis approaches to predict and understand drug-protein binding specificity using imatinib binding to Abl, Chk1, Kit, Lck, p38α, and Src as model systems. All co-crystal structures of imatinib bound to kinases show that the drug fits well within the binding pocket, except for imatinib-Chk1, for which no experimentally determined structure is available. Multiple MD runs were performed for each system, including both imatinib-bound and free-state kinases. Subsequent analysis of the MD trajectories yields quantitative thermodynamic components of imatinib-kinase binding, revealing key contributions such as enthalpy and dihedral configurational entropy. Force distribution analysis (FDA), principal component analysis (PCA), and sidechain dihedral correlation networks were used to assess protein dynamics effects on drug binding specificity. This study shows that thorough analysis of drug-bound MD simulations accurately identifies molecular determinants of binding specificity. These findings may inform RDD by revealing distinct conformational changes associated with imatinib’s affinity for its target kinase.

## Method

### Molecular Systems

The initial structures of Abl, Kit, Lck, p38α, and Src were the protein complex crystal structures with imatinib in the ATP-binding site (PDB ID: 2HYY for Abl, PDB ID: 1T46 for Kit, PDB ID: 2PL0 for Lck, PDB ID: 3HEC for p38α, and PBB ID: 2OIQ for Src).^20,22,31–33^ The initial structures of Chk1 were the protein complex crystal structures with 6-substituted indolylquinolinones in the ATP-binding site (PDB ID: 2HXL) in the DFG-in active configuration.^34^ Previous systematic reviews affirm that Chk1 kinase is only found in the DFG-in form.^35^ Therefore, Chk1 is hypothesized to bind with imatinib in its DFG-in form. While imatinib typically binds in the DFG-out inactive structure, it is also observed to bind with proteins in a compact cis form, including imatinib-bound Syk kinase and Noq2 reductase (Figure 2).^36–38^ The imatinib-bound Syk kinase (PDB ID: 1XBB) was used to obtain a reasonable conformation for bound imatinib for superimposition, and copy-paste alignment was used to replace the 6-substituted indolylquinolinone in Chk1.^23^ Imatinib adopts two protonation states in solution, with previous experiments indicating that it binds to TKs in the monoprotonated form.^28,39–42^ Therefore, the monoprotonated form was used for all experimental proteins and bulk solution runs. The chain A protein and crystal water molecules were captured for all systems. Abl was missing Glu275, and SWISS-MODEL was used to correct the structure.^43^ In the imatinib-bound Kit complex, the missing residues from Ile690 to Glu761 in a distal loop that substituted the kinase insert domain were transplanted from sunitinib-bound Kit (PDB ID: 3G0E) to fill the missing loops.^44^ The imatinib-bound p38α kinase was missing two loop regions from Ser32 to Gly36 in the phosphorylation loop (P-loop) and Gly170 to Val183 in the activation loop (A-loop) following the DFG motif. A reasonable conformation using another DFG-out p38α complex bound to a small molecular inhibitor (PDB ID: 1W82) was used to fill the structure with copy-paste alignment.^45^ Src kinase was missing the A-loop from Leu407 to Gly421. A reasonable configuration was inserted with copy-paste alignment and mutated with Molecular Operating Environment using imatinib-bound Lck as a reference.^33^ To consistently preserve all key interactions observed in the crystal structures, all crystal waters in the original PDB were retained.

**Figure 2.**
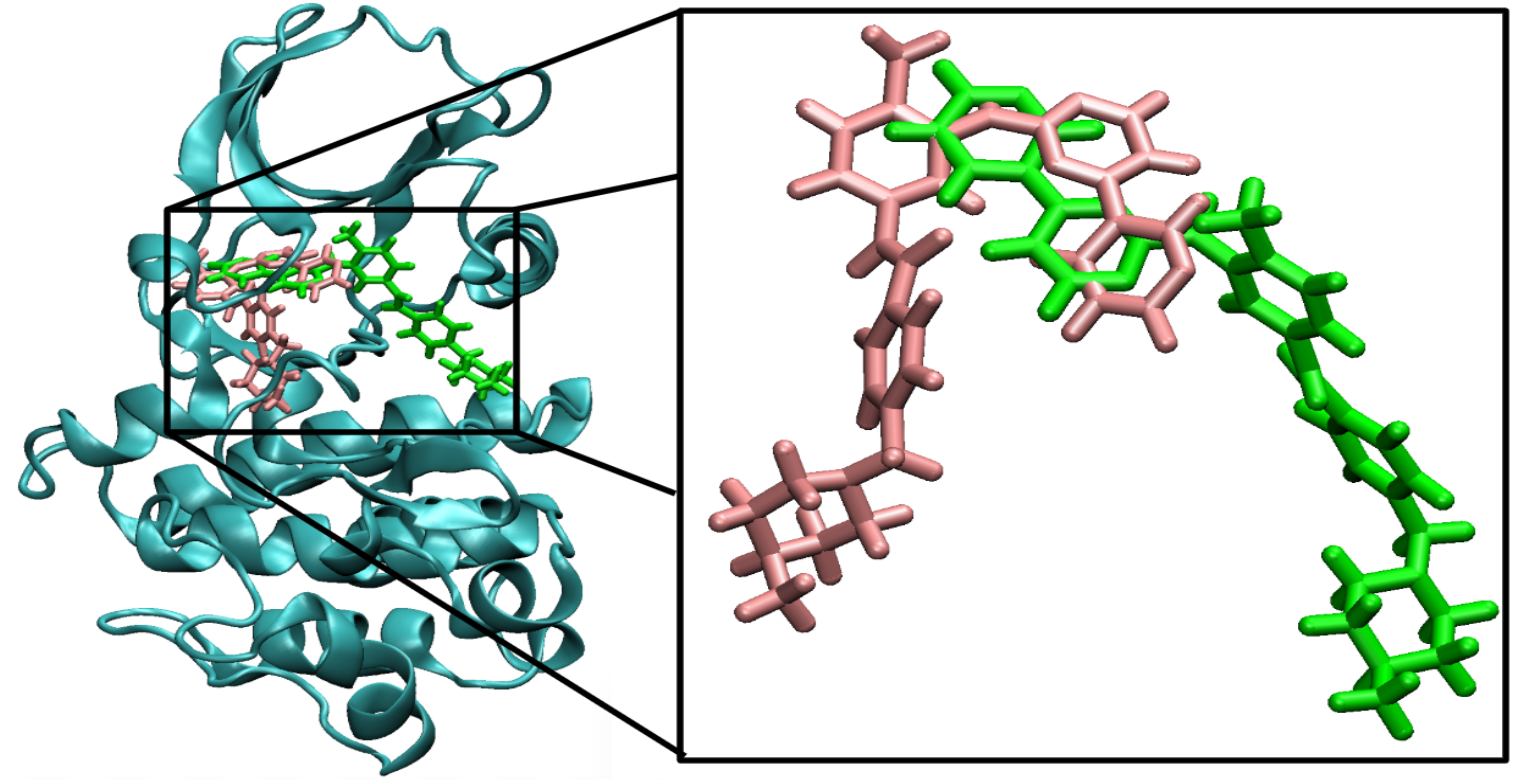
3D structure of Chk1-bound cis (pink) and Abl-bound trans (green) imatinib configuration showing alternate binding modes.

### Simulations Protocol

The MD simulations were run using the Amber package.^46^ All MD runs used Amber ff14SB for the proteins, Generalized Amber Force Field (gaff2) for imatinib, and tip3p waters for the solvent.^47–49^ The tleap module was used to add all missing hydrogens with special attention paid to histidine delta and epsilon protonation states based on the local environment.^50^ PROPKA was used to verify the protonation states of residues.^51^ Imatinib was assigned AM1-BCC charges with the antechamber module in Amber.^52^

The hydrogen atoms, amino acid sidechain, and whole protein system were minimized for 500, 1000, and 5000 steps, respectively, in a generalized Born implicit solvent. A cuboid water box was used to attain the same amount of water molecules, 22,307, for a standardized amount. This amount ensures that all systems in this study were solvated in a water box with a distance of at least 12 Å from the edge of a kinase. The Parmed software in Amber was used to adjust the number of water molecules to enable binding enthalpy calculations. One chloride counterion was added to counterbalance the imatinib’s +1 charge state.^28,39–42^ The remaining systems were solvated with the appropriate amount of sodium counterions to achieve a neutral state. This was 8, 0, 7, 7, 5, and 6 for Abl, Kit, Lck, p38α, Src, and Chk1 kinases, respectively, at the physiological salt concentration of 150 mM. Free charged imatinib was run in solution with one chloride ion and a cuboid box filled with 22,307 waters to provide the enthalpy calculations and compensate for an extra missing water box. The hydrogen-built systems for solvated and neutralized imatinib-bound systems consist of 71,265, 71,457, 71,919, 71,352, 72,610, and 71,455 atoms for the Abl, Chk1, Kit, Lck, p38α, and Src systems, respectively.

The water molecules were minimized for 10000 steps, followed by minimization of the whole system for 20000 steps. The solvated system was equilibrated under constant pressure and temperature (NPT) ensemble from 50 K to 275 K with 25 K increments and 200 ps each, and finally at 300 K for 500 ps. Langevin thermostat and Berendsen barostat were used for an isothermal and isobaric condition at 298 K and 1 Bar. The SHAKE algorithm was used to constrain all bonds with hydrogen atoms.^53^ 2 femtosecond timesteps were used for the simulations. Long-range electrostatic interactions were considered using the particle mesh Ewald scheme, with a nonbonded cutoff of 12 Å. The simulations were conducted using an NPT ensemble with an implicit water energy minimization algorithm. All simulations were run for 500 ns using a timestep of 2 fs at 298 K for all proteins, free imatinib, and the water box. The protein system may get trapped at one local minimum, yielding suboptimal results. Three repeats for each bound-protein, free protein, free imatinib, and water box were performed for varied sampling. The first 50 ns were treated as equilibration and discarded, leaving 450 ns for the post-analysis.

### Interaction Energy Calculation

A frame was resaved every 500 ps through a 500-ns MD trajectory to generate 1000 frames for Molecular Mechanics Poisson-Boltzmann surface area (MM/PBSA) energy calculations. Amber MM/PBSA was used to estimate the relative interaction energy of a complex system consisting of imatinib and a kinase.^54^ For a single trajectory, the energy terms were separated for the complex, receptor, and ligand. The relative changes are calculated using the formula: *ΔE = E*_*complex*_ - *E*_*receptor*_ *-E*_*ligand*_. All calculations utilized an external dielectric constant of 15.0.

### Binding Free Energy Calculations

#### Enthalpy Calculations

Custom scripts in R were used to extract total energy (E_tot_) values for the MD ensembles from each output file every 1000 ps throughout a 500-ns MD trajectory for 500 points. The mean convergence was calculated after the main trajectory equilibrated to approximate and visualize enthalpy using the dplyr and ggplot2 packages in R language. The enthalpy range was calculated based on the upper and lower bounds to encapsulate all combinations of complex, receptor, ligand, and water box across repeats. The formula used is *ΔH = E*_*tot,complex*_ *– E*_*tot,receptor*_ *– E*_*tot,ligand*_ *+ E*_*tot,waterbox*,_ which compensates for a missing water box in the total system. The upper-bound *ΔH* is approximated with *ΔH*_*upper*_ *= E*_*tot_complex_minimum*_ *– E*_*tot_receptor_maximum*_ *– E*_*tot_ligand_maximum*_ *+ E*_*tot_water_minimum*_. The lower-bound *ΔH* is approximated with *ΔH*_*lower*_ *= E*_*tot_complex_maximum*_ *– E*_*tot_receptor_minimum*_ *– E*_*tot_ligand_minimum*_ *+ E*_*tot_water_maximum*_. This novel method provides the minimal and maximal enthalpy ranges for all combinations of the independent simulations, offering a comprehensive overview of the bound-kinase, free kinase, and free imatinib thermodynamics.

#### Dihedral Entropy Calculations

The T-Analyst program was used to calculate the changes of dihedral entropies in the protein binding site and rotatable bonds of imatinib for the imatinib-bound complex, free protein, and free imatinib.^55^ A frame was resaved every 50 ps through a 500-ns MD trajectory to generate 10000 frames for dihedral entropy analysis. T-Analyst applies the Gibbs entropy formula −*RT ∫ ρ*(*x*)*logρ*(*x*)*dx* ≈ −*RT* ∑ *ρ*(*x*)*logρ*(*x*)*dx* to compute dihedral entropy with numerical methods, where R is the ideal gas constant, T is the absolute temperature (298K), *ρ*(*x*) is the probability distribution, and dx is the bin width.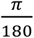 radians (1º) were used for phi and psi angles, and 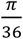 radians (5º) were used for sidechains. Histograms are used to categorize the probability distribution of the phi, psi, and sidechain dihedrals. The Gibbs entropy formula can numerically yield a negative value if the distribution has a sharp unimodal peak. These negative values are not considered as they are not physically meaningful. The Boltzmann equation S = −*kBln* (Ω) relates entropy with kB, the Boltzmann constant, and Ω the number of accessible microstates. The condition where the torsion angle distribution has a sharp unimodal peak corresponds to one microstate, such that S = −*RTln* (1) = 0. Calculating the changes in entropy uses the formula: *ΔS = S*_*complex*_ *-S*_*receptor*_ *-S*_*ligand*_. An upper and lower bound was calculated based on the output. The lower-bound entropy is approximated with *ΔS*_*lower*_ *= S*_*complex_minimum*_ *-S*_*receptor_maximum*_ *-S*_*ligand_maximum*_. The upper-bound entropy is approximated with *ΔS*_*upper*_ *= S*_*complex_maximum*_ *-S*_*receptor_minimum*_ *-S*_*ligand_minimum*._ This innovative approach gives the minimal and maximal entropy range for all combinations of the independent triplicate simulations.

These enthalpy and entropy calculations were used to calculate the changes in Gibbs free energy using the *ΔG = ΔH* - *TΔS* formula. The upper-bound Gibbs energy is calculated with *ΔG*_*lower*_ *= ΔH*_*lower*_ – *TΔS*_*upper*_. The lower-bound Gibbs energy is calculated with *ΔG*_*upper*_ *= ΔH*_*upper*_ – *TΔS*_*lower*_. These results capture an upper and lower bound that encompasses all independent combinations for the complex, receptor, ligand, and water box, providing an innovative and comprehensive overview of the kinase and imatinib binding thermodynamics for the triplicate ensembles.

### Force Distribution Calculations

Force distribution analysis (FDA) implemented in GROMACS is used to attain atomistic insights into force vector magnitude and directionality for protein residues.^56^ This gives novel insights into the locomotion of key secondary structures for the DFG-in and DFG-out binding modes. A frame was resaved every 50 ps through a 500-ns MD trajectory to generate 10000 frames for force calculations. A cutoff of approximately ±5 pN was used to disregard residues forming weak interactions. From the data collected, the residues are identified as constituting the binding pocket across all six protein-drug complex sets and three experimental repeats for a robust collection. The FDA reveals protein residues that form important and underexplored non-covalent interactions with imatinib, including those near β-sheets, the catalytic loop, and the αF-helix, thereby enhancing the data set’s accuracy and novelty.

### Residue Sidechain Correlation

Residue sidechain correlation analysis is used to capture important coordinated dihedral motions in the MD simulations. The T-Analyst program was used to calculate the sidechain dihedral rotations of all amino acid residues and their pairwise correlations with every other residue sidechain in the proteins using the Gibbs entropy formula.^55^ A frame was resaved every 50 ps through a 500-ns MD trajectory, providing 10000 frames for torsion angle calculations. Pairwise correlations are calculated using a Pearson correlation formula. A correlation matrix is plotted. Cartesian coordinates are transformed to bond-angle-torsion coordinates to capture the differences and prevent unusual discontinuity margins close to ±π radians (±180 degrees) or 2π/0 radians (360/0 degrees). Positive correlation corresponds to two sidechains moving similarly, negative correlation means they move in the opposite direction, and a correlation close to 0 means no coordinated motion. From the pairwise correlation matrix, a correlation network is generated using the Python library NetworkX to visualize the residue-residue correlation links across the three experimental ensembles.^57^ A cutoff of ±0.30 for Abl, Lck, p38α, and Src; ±0.25 for Chk1, and ±0.20 for Kit is applied to eliminate residues with negligible correlated motions. The residues were matched with their corresponding secondary structure and marked based on their known function in the protein. If a residue-residue correlation beyond the threshold was observed for two or more repeats, it was noted as a significant link and included for 3D network visualization. Visual Molecular Dynamics (VMD) was used to visualize the residue-residue correlation by placing links between the α-carbons across the experimental triplicates.^58^

### Principal Component Analysis

Principal component analysis (PCA) is used to elucidate essential dynamics of a protein system by reducing data dimensionality to the top principal components (PCs). In this study, the BKiT package is used to calculate essential α-carbon PCA motions for each bound-complex and apo-state MD run.^59^ A frame was resaved every 50 ps through a 500-ns MD trajectory to generate 10000 frames for PCA calculations. PCA characterizes the variance magnitude of individual residues and the whole secondary motifs of the P-loop, αC-helix, and A-loop to show their migration patterns in 3D Cartesian space. The top contributing motions, identified with the eigenvectors, indicate the general migration pattern of key regulatory motifs. These migration patterns reveal if the protein adopts a more open or closed configuration on average, enabling ATP competition with the imatinib in the binding pocket. The formula for PC mode calculation is *qi = R*^*T*^ *(x*_*i*_ *-x*_*a*_), where x represents the conformation along the trajectory, i represents the principal component on one PC mode, and R^T^ is the transposition of the eigenvector matrix of the covariance matrix. VMD was used to visualize the magnitude of the first PC across ensembles for general insights into secondary structure movement and to identify which residues contributed the greatest captured variance across sets.^58^

## Results & Discussion

### Computing Interaction and Binding Free Energy Using MD Trajectories

MD simulations are a widely used method to simulate the physical movements of proteins and their complex with a bound inhibitor. Beyond sampling molecular conformations, one can extract protein-ligand binding enthalpy, entropy, and free energies. One can further dissect the molecular determinants of each thermodynamic component by post-processing MD trajectories. The computed binding free energy (ΔG), as well as the relative binding free energy (ΔΔG), can be directly compared with experimental measurements. Notably, while the trend in imatinib’s binding affinity across different kinases remains consistent, the absolute values vary depending on the experimental conditions. Tables S1 and S2 summarize the experimental binding and inhibition data, K_D_ and IC_50_, for imatinib across the six kinases examined in this study.

MM/PBSA energy calculations were performed on the MD trajectories of the imatinib-kinase complexes to estimate their binding affinity. As reported in Table S1 and Figure S1, the computed interaction energies do not accurately reproduce the experimental results. While the calculation is a widely used post-analysis strategy, the results highlight the limitations of relying solely on intermolecular interactions to capture binding. While the MM/PBSA calculations can reasonably quantify non-covalent intermolecular interactions and solvation free energy changes upon binding, the entropic contribution is neglected. Moreover, because only the imatinib-kinase MD trajectories are used, conformational free energy changes are not accurately captured.

The post-processed MD trajectories for the imatinib-kinase bound complexes, free kinases, free imatinib, and a water box were used to estimate the binding free energies more accurately. The same number of water molecules was maintained across all MD runs to apply a direct method for estimating binding enthalpies, *ΔH = E*_*tot,complex*_ *– E*_*tot,receptor*_ *– E*_*tot,ligand*_ *+ E*_*tot,waterbox*,_, where the total energy *E*_*tot*_ was extracted from the MD output.^47^ Entropy changes upon imatinib binding were approximated by computing dihedral entropy for the bound complex, free kinases, and free imatinib, as detailed in the Method section. Because each independent MD run sampled a subset of the full conformational ensemble, for each imatinib-kinase system, the upper and lower bounds of the corresponding energy and dihedral entropy were approximated based on the computed maximum and minimum values. The range of computed ΔG is estimated using the difference between the maximum enthalpy change (ΔH_upper_) and the minimum entropy change (TΔS_lower_), as well as the difference between the minimum enthalpy change (ΔH_lower_) and the maximum entropy change (TΔS_upper_), shown in Table S4. Table 1 and Figure 3 present the ranges of computed and experimental ΔG, as well as experimental IC_50_ values, with individual experimental data points and corresponding references detailed in Tables S1 and S2. All ΔG_comp_, ΔG_exp,_ and IC_50,exp_ consistently show that imatinib binds most tightly to Abl and Kit, with affinities in the low nanomolar range; binds modestly to Lck and Chk1 in the low micromolar range; and exhibits the weakest binding to Src and p38α in the high micromolar range. These results show that explicit MD simulations can be effectively exploited to accurately capture relative changes in ΔG_comp_.

**Table 1.**
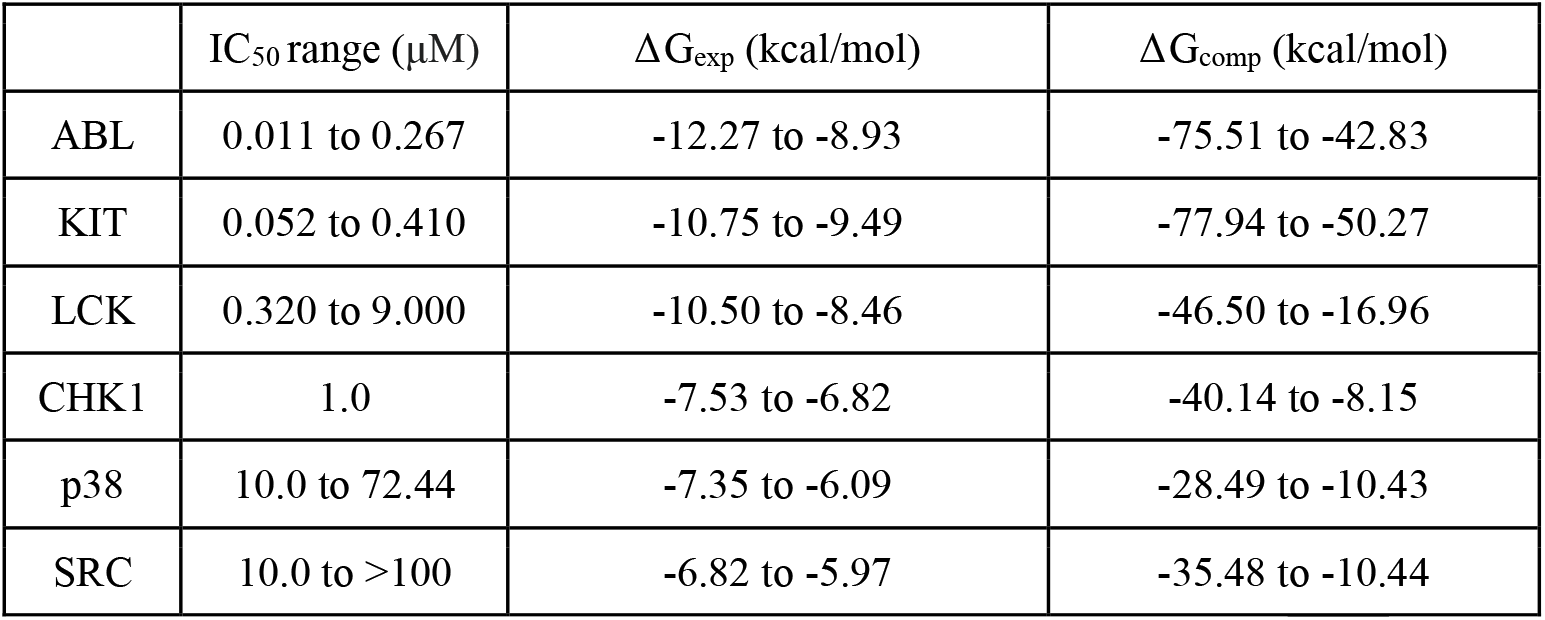
Imatinib IC_50_ inhibition, experimentally measured binding affinity ΔG_exp_, and computed binding affinity ΔG_comp_ from enthalpy and entropy calculations using MD trajectories. The collection of the experimental data is found in the SI.

**Figure 3:**
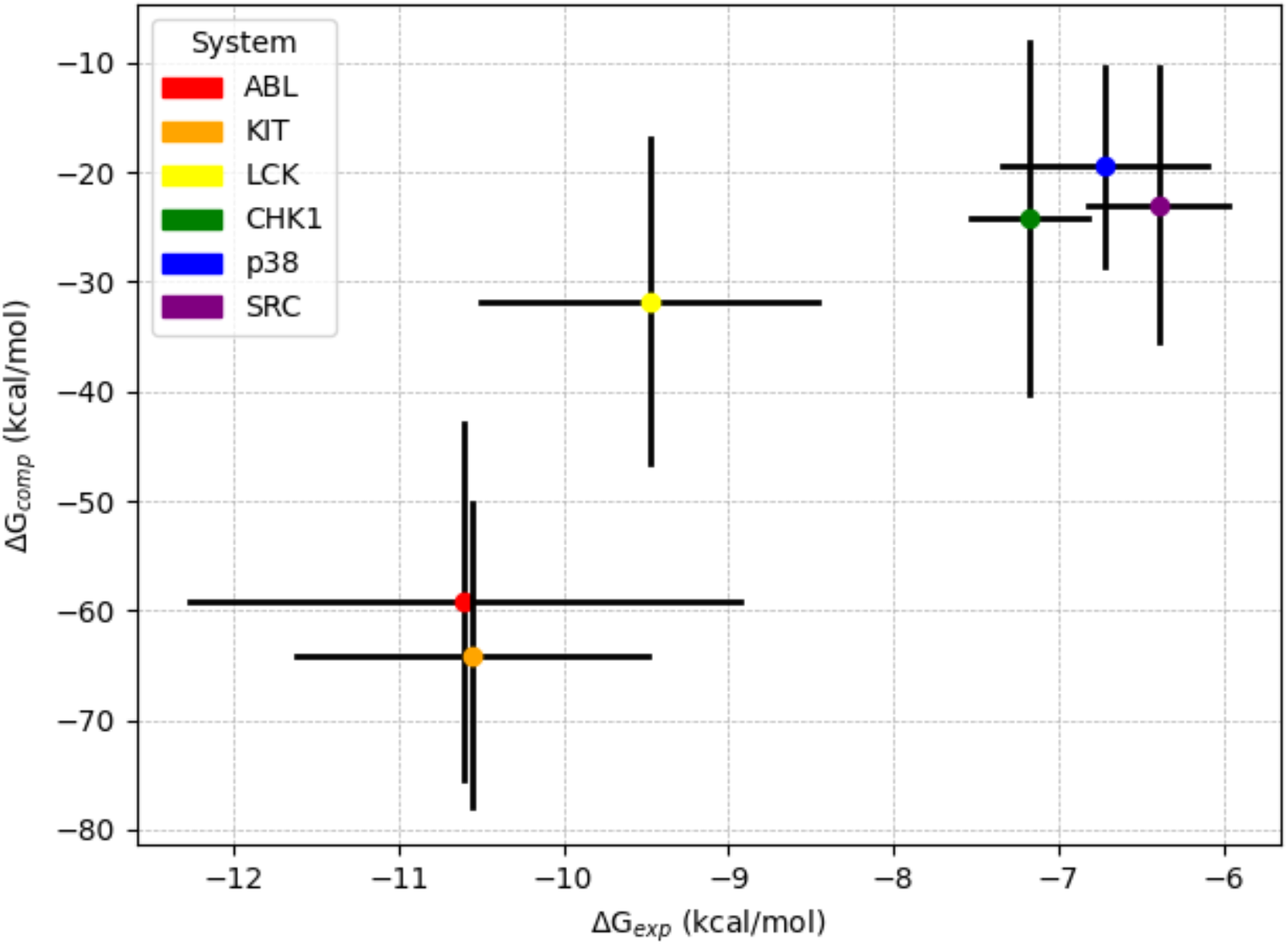
Comparison of experimental and computed binding free energy values. The colored dot is the median value of the possible range, while the black bars are the lower- and upper-bounds of the range. The Pearson R correlation is 0.92.

### Assessing Force Vector Directionality with Force Distribution Analysis

Structural dynamics and molecular interactions provide clues for imatinib-kinase specificity insights by clearly identifying the key residues implicated in binding. Root-mean-square deviation and fluctuation (RMSD and RMSF) analysis of the α-carbons shows that all the simulations reach an equilibrated state, and which regions are more flexible (Figures S3 and S4). But neither can show which directions fluctuations occur, missing critical insights for specific configurational dynamics. FDA accounts for this by calculating the average force vector for the protein residues and imatinib atoms in all 10000 frames across the trajectory to give directional insights for kinase motif locomotion. A key advantage is that it characterizes these motions more accurately than just the attractive or repulsive energy terms with an interaction energy calculation method like MM-PBSA. A cutoff of approximately ±5 pN for either attraction or repulsion is used to accurately and comprehensively select for the binding site residues with reasonably strong non-covalent imatinib interactions (Figures S5 to S10).^60^ The observed residues forming a strong interaction are generally found in key regulatory motifs, such as the P-loop, the αC-helix, and the A-loop, along with other underexplored regions, including the β3 sheet, catalytic loop, hinge region, αE-helix, αF-helix, and connector loops. They are observed across multiple repeats, demonstrating the robustness of the force vector method to characterize the pocket residues and show the direction of motif motions. The findings corroborate experimental data showing that mutations in these proximal regions disturb imatinib binding specificity in Abl kinase.^61–66^ These findings likewise agree with observed mutations in regions disturbing Kit kinase sensitivity to imatinib.^44,64,67–69^

The FDA results show the same general regions form strong interactions; particularly the P-loop, αC-helix, hinge region, and A-loop (Figures S5 to S10). Abl kinase tends to strongly interact with the P-loop, the hinge region, and the A-loop, and form weaker interactions with the αC-helix (Figures 4 and S5). KIT kinase forms weaker interactions with the P-loop across the runs, while forming stronger interactions with the αC-helix, hinge region, and A-loop (Figure S6). The Chk1 kinase has a distinct binding mode and tends to form weaker interactions with the P-loop and stronger interactions with the αC-helix, hinge region, and A-loop (Figures 5 and S7). Lck kinase forms stronger interactions with the P-loop and hinge region, and fewer interactions with the αC-helix and A-loop (Figure S8). The p38α kinase typically interacts with the αC-helix, hinge region, and A-loop and interacts less with the P-loop (Figure S9). The Src kinase forms stronger interactions with the αC-helix, hinge region, and A-loop and has fewer interactions with the P-loop (Figure S10).

**Figure 4.**
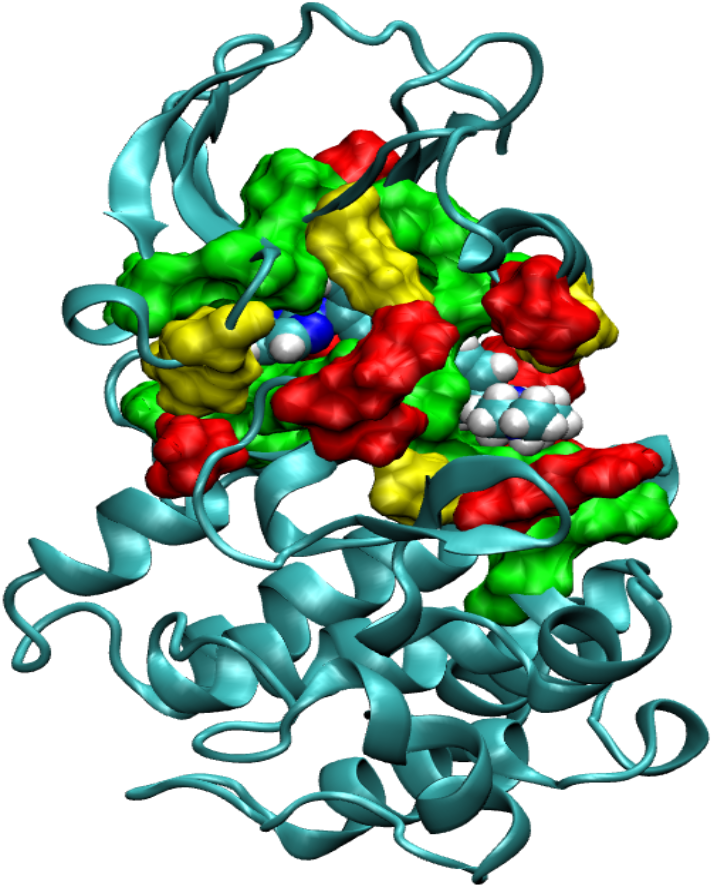
ABL1 kinase and residues with strong force interactions. Residues exhibiting ∼ ±5 pN force on average shown in surface. Green, yellow, and red exhibit on average more than ±5 pN of force in three, two, and one MD runs, respectively. The snapshots are captured at the end of a 500 ns trajectory.

**Figure 5.**
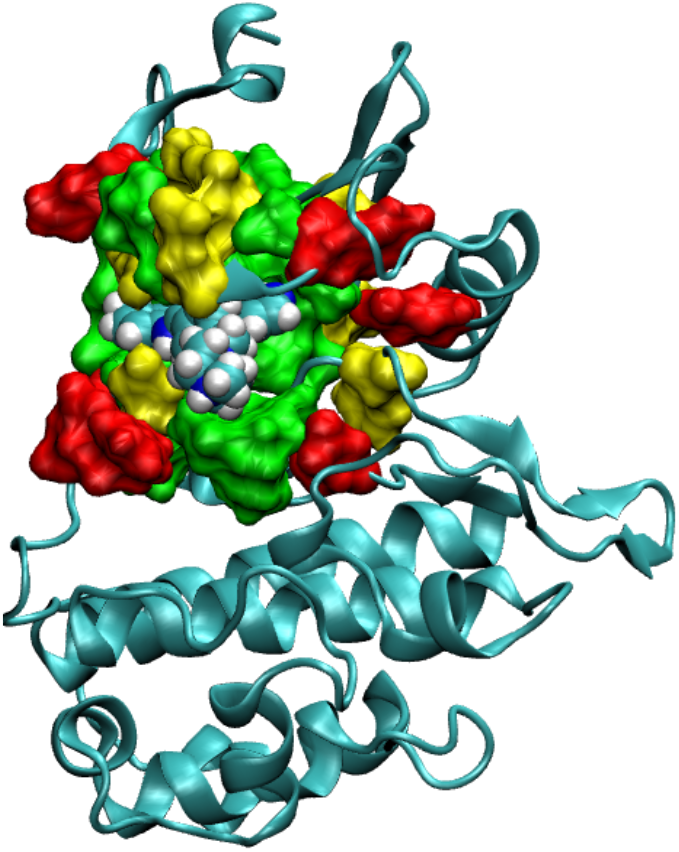
CHK1 kinase and residues with strong force interactions. Residues exhibiting ∼ ±5 pN force on average shown in surface. Green, yellow, and red exhibit on average more than ±5 pN of force in three, two, and one MD runs, respectively. The snapshots are captured at the end of a 500 ns trajectory.

The observed imatinib-bound Abl residues interactions are consistent with previous collections of experimental data as Leu248, Tyr254, and Val256 in the P-loop; Glu282, Glu286, Val280, Met290, and Ile293 in the αC-helix; Ile313 to Asn322 encompassing the hinge region; and Asp381, Phe382, Ser385, and Arg386 in the A-loop, all significantly influence the Abl binding pocket domain affect the inhibition capacity of imatinib for the mutants (Figure 1A). Mutations in these nearby regions are implicated in reduced imatinib sensitivity, such as the Thr315 gatekeeper mutant to Ile.^65,70,71^ The observed imatinib-bound Kit residues are also consistent with prior reported literature, where Leu595, Val603, Val604, and Glu605 in the P-loop; Glu640, Val643, Leu644, and Leu647 in the αC-helix, Val668 to Asp677 in the hinge region; and Ile808 to Phe811 in the A-loop. Mutations in these proximal residues correspond with reduced patient responses and increased remission when taking imatinib, such as the Thr670 gatekeeper mutant to Ile.^67–69^ These insights show that, even though the binding pocket is highly similar across proteins, there are observable differences that influence specificity in the higher-affinity Abl and Kit kinases and the corresponding resistance mutation changes disrupting imatinib sensitivities.

### Discerning Secondary Motif Migration in Cartesian Space with Principal Component Analysis

PCA is a tool to identify important kinase structural migration patterns after imatinib binding, using the movements of the α-carbons in the key structural motifs. The largest protein motions are usually the most biologically relevant and effectively predict binding site exposure to solvents and competitive ligands, like ATP.^72^ After imatinib binds, the Abl and Chk1 PCA shows the P-loop and αC-helix shift towards the pocket, and the A-loop moves in a way consistent with adopting the inactive kinase configuration (Figures S11 and S12). The Kit and Lck kinases show less pronounced P-loop motions, the αC-helix closes into the pocket, and the A-loop broadly adopts an inactive form (Figures S13 and S14). These findings show that in higher-affinity proteins such as Abl, Kit, Lck, and Chk1 kinase, the essential motions of the regulatory motifs tend to restrict solvent and ligand access to the ATP pocket (Figures 6 and S11 to S14). In the p38α and Src kinases, the broad movements show that the αC-helix moves the most towards an active pocket-exposed configuration, while the A-loop also moves in a way to adopt an active configuration in Src (Figures 7, S15, and S16). These trends show that in the lower-affinity proteins like p38α and Src, there is a tendency for the structure to open, increasing solvent and ligand exposure to the pocket. These reports are consistent with previously reported solvent accessible surface area insights from crystal structures.^22^ If the kinase secondary motifs modulate access to the pocket and tend to close in the higher-affinity proteins and open more regularly in the lower-affinity kinases, then the solvent and competing molecules may access the protein interior more readily.

**Figure 6.**
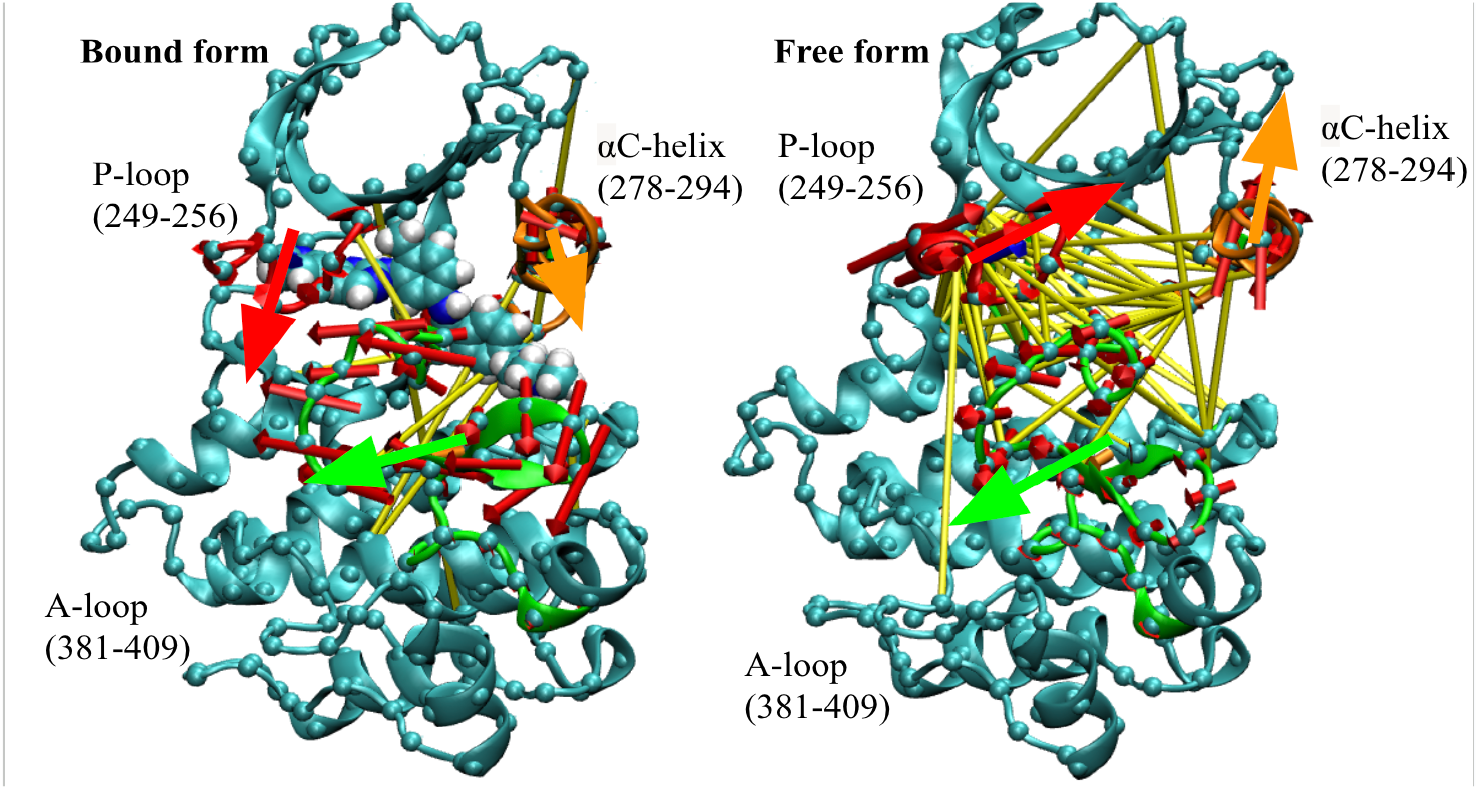
ABL sidechain correlation network and breathing motions of the P-loop, αC-helix, and A-loop triad identified by α-carbon PCA. The yellow and green cylinders correspond with an (anti-)correlation cutoff of ±0.3 for the residue sidechains in two and three experimental repeats. The P-loop motions are shown in red, the αC-helix in orange, and the A-loop in green. The large red, orange, and green arrows show the average migration pattern for the secondary motif based on the eigenvectors for the first PC calculated from BKiT. The snapshots are captured at the end of a 500 ns trajectory.

**Figure 7.**
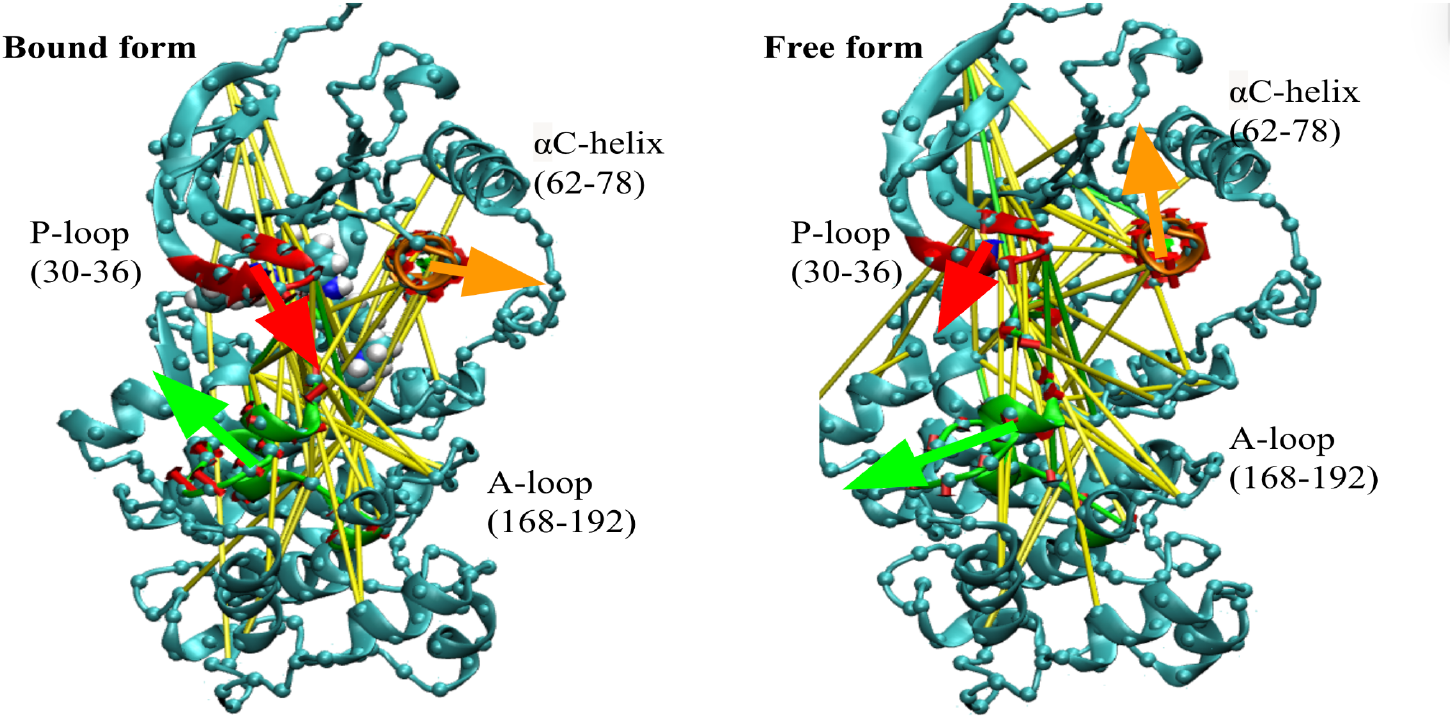
p38α sidechain correlation network and breathing motions of the P-loop, αC-helix, and A-loop triad identified by α-carbon PCA. The yellow and green cylinders correspond with an (anti-)correlation cutoff of ±0.3 for the residue sidechains in two and three experimental repeats, respectively. The P-loop motions are shown in red, the αC-helix in orange, and the A-loop in green. The large red, orange, and green arrows show the average migration pattern for the secondary motif based on the eigenvectors for the first PC calculated from BKiT. The snapshots are captured at the end of a 500 ns trajectory.

### Revealing Essential Protein Conformational Network with Sidechain Dihedral Correlations

Dihedral angles are the major variables to govern conformational changes, analyzing their correlations provides a more targeted analysis of functionally relevant motions, particularly for the P-loop, αC-helix, and A-loop.^73^ They are the degrees of freedom enabling the kinase to adopt distinct configurations, and are more accurate compared to use of Cartesian coordinates since they inform whether the structures are changing in a coordinated way. We used different cutoffs based on the kinase’s observed rigidity, as the sensitivities for coordinated protein motions are highly system-dependent due to differing amino sequences, folding topology, and conformational landscaping. The findings generally show that the sidechains making up the key regulatory motifs of the P-loop, αC-helix, and A-loop move in coordinated manners across MD runs for the six free-form proteins. For stronger-binding proteins such as Abl, Kit, and Lck, there is an overall tendency towards network collapse as the regulatory motifs exhibit reduced communication and fewer appropriate configurations for ATP competition after imatinib binds. Chk1 shows a unique deviation, where imatinib binding corresponds with a stronger bound-structure dihedral network. As imatinib is hypothesized to bind Chk1 in the active form, this indicates that facilitating more dynamic movements may disturb the active form structure and reduce ATP competition for the pocket-bound imatinib. In contrast, weaker-binding proteins like p38α and Src tend to retain or enhance the existing network formation for those key regulatory motifs, enhancing communication across the protein structure to adopt configurations more likely to have increased ATP competition. This suggests that modulating the fidelity of information transmitted across a protein structure is a key imatinib specificity factor.

The residue sidechain correlation data also identifies distal mutants further from the binding site. In Abl, more distal residues captured included Thr277 in the β3-αC loop, Arg307 in the β4-β5 loop, Phe359 in the catalytic loop, Val371 in the β7 motif making up the hydrophobic cage, and Ile418 and Leu429 in the αF-helix across at least two independent repeats. Mutants at or near these distal sites, including Thr277, Phe359, Val371, and Ile418, are observed to experience weakened imatinib sensitivity.^65^ Likewise, for the Kit systems, key distal residues found using the correlation network include Ile653 in the αC-β4 loop and Arg815, Asp816, and Met836 in the A-loop. These regions form key interactions influencing the protein’s structure and host mutations that deleteriously affect imatinib’s ability to bind and inhibit Kit, including Val654, Asp816, and Arg830.^44,68,74–76^ This shows the predictive power of the dihedral correlation network to identify deleterious mutants via a hidden conformational network, corroborating previous drug binding and mutation studies on distal regulatory communication in these kinases.^77–79^

The structure-function relation for proteins necessitates that coordinated dynamics, rather than random motions, must guide a protein’s structural arrangement to perform its necessary physiological actions. As an example, kinases adopt a “breathing motion” where the regulatory motifs of the P-loop, αC-helix, and A-loop move in a coordinated way to let ATP access and leave the pocket.^80–83^ These motions are broadly seen using force vectors from FDA and α-carbon essential motions from PCA. More specific insights are elucidated using internal torsion dynamics to filter for functional motions. The correlated sidechain motions were also investigated to discern the functionally relevant motions of residues that move together in the structure. The above analysis reveals that FDA residue identification, sidechain dihedral correlation analysis, and α-carbon Cartesian PCA reveal a hidden configurational network through protein residue sidechains. This network is observed consistently across experimental triplicates, and imatinib’s effect on the network’s ability to transmit structural data through key residue nodes is clearly visible (Figures. 6 and 7). Conformational data is transmitted across the protein through these factors, and the strength of the network corresponds with the fidelity of the necessary structural change. The combined network and essential motion analysis shows that in higher-affinity proteins such as Abl, Kit, and Lck kinases, there is a tendency towards network collapse across multiple system ensembles, disrupting essential protein configurational communication networks, while this network is strengthened in the lower-affinity p38α and Src kinases.

This hidden dihedral network allows kinases to perform their key function of binding with ATP to facilitate phosphate transfer to natural substrates on other proteins for cell signaling. The network correlation and essential dynamics data show that when imatinib binds favorable proteins like Abl, Kit, and Lck, it disrupts this coordinated breathing motion, limiting the proteins’ ability to adopt a favorable configuration for ATP approach and competition across ensembles. Chk1 also shows a disruption in the breathing motion from Cartesian PCA, but a strengthened communication network, suggesting a different mechanism corresponding with the alternative binding mode. When imatinib binds to less favorable proteins like p38α and Src, it retains most of the existing connections, prompting the ATP pocket to open by maintaining coordinated motions with other key regulatory motifs. This corresponds with the protein being more likely to adopt an open configuration and promoting ATP competition. This will increase the likelihood of imatinib dissociation across ensembles and reduce its observed affinity, reflected in both our computed binding affinity ΔG_comp_ from our enthalpy and entropy approximations and experimentally measured ΔG_exp_. This study shows the importance of the kinase regulatory motifs to adopt a favorable configuration and does so through coordinated motions not previously observed in literatures studying these kinases’ structural or dynamic properties. Kinase loops between the regulatory motifs also play an important role in coordinating information across the protein in ways not previously characterized.

## Conclusions

In this study, we present strategies to systematically investigate and elucidate drug promiscuity with multiple kinases using imatinib as a model system. Rigorous testing with four TKs (Abl, Kit, Lck, and Src) and two STKs (Chk1 and p38α) demonstrates the approach to use thermodynamics calculations, FDA, residue correlation networks, and PCA essential motions to explain kinase-imatinib specificity across multiple kinases. These factors reveal a hidden network spreading information about the preferred kinase configuration following imatinib binding. The reported dynamics data show that strong and intermediate imatinib-binding kinases, such as Abl, Chk1, Kit, and Lck, undergo disrupted ATP competition due to interrupted bound-kinase breathing motions, whereas the weaker imatinib-binding proteins, like p38α and Src, enhance ATP competition due to promoted bound-kinase breathing motions. These findings may have strong implications for future RDD to construct effective binders that selectively interrupt kinase domain communication networks. Future work to further characterize and dissect the roles of coordinated motions on drug binding will be pursued.

## Supporting information

Troxel_Supporting_Information

## Data Availability

The molecular structures are displayed in VMD (https://www.ks.uiuc.edu/Research/vmd/). The GROMACS program is used to calculate the force vector interaction between the imatinib and protein kinase residues (https://www.gromacs.org/). The in-house BKiT program is used to perform principal components analysis (https://github.com/chang-group/BKiT). The in-house T-Analyst program is used to calculate sidechain configurational entropy and dihedral correlations (https://www.chang-group.org/software/). The supporting topology files, trajectories, calculation scripts, input files, and output files are openly available upon request.

## Acknowledgements

Research support was provided by the US National Institutes of Health (T32ES018827 and GM109045), the US National Science Foundation (MCB2437134), and the Department of Education (K012825001). Computations were performed using the high-performance computer cluster, ACCESS, from the NSF (CHE130009) and UCR High Performance Computer Cluster. We thank Dr. Jianan Sun to assist with the BKiT package.

