## Supplementary material for "Revealing imatinib-kinase specificity via analyzing changes in protein dynamics and computing molecular binding affinity": Troxel_Supporting_Information

William Troxel[1], Emily Vig[2], Chia-en Chang[1,2]*
1. Department of Chemistry, University of California, Riverside, California 92521, United States
2. Department of Biochemistry and Molecular Biology, University of California, Riverside, California 92521, United States

*Correspondences
Chia-en Chang:

**This PDF file includes**

Supplementary Text
Figs S1 to S16
Tables S1 to S4
References

Table S1. Mean convergence of MM/PBSA interaction energy analysis for protein-drug complex for a 500 ns molecular dynamics (MD) trajectory (n = 3).

| | MD run 1 | MD run 2 | MD run 3 |
|---|---|---|---|
| **ABL** | -39.19831489 | -36.54394811 | -35.96928067 |
| **CHK1** | -23.49748411 | -23.52476133 | -25.44881956 |
| **KIT** | -36.87133622 | -36.57166033 | -35.62990367 |
| **LCK** | -35.32584233 | -36.42021589 | -35.81523889 |
| **p38α** | -32.678528 | -36.695207 | -32.154196 |
| **SRC** | -40.279773 | -36.04763656 | -37.57990967 |

Table S2. Enzymatic inhibitory assays of imatinib across multiple kinases

| | $IC_{50}$ (μM) | Reference Paper |
|---|---|---|
| ABL | 0.0108 | [1] |
| ABL | 0.011 | [2] |
| ABL | 0.025 | [3] |
| ABL | 0.025 | [4] |
| ABL | 0.025 to 0.200 | [5] |
| ABL | 0.025 to 0.200 | [6] |
| ABL | 0.025 to 0.200 | [7] |
| ABL | 0.025 to 0.200 | [8] |
| ABL | 0.037 | [9] |
| ABL | 0.038 | [8] |
| ABL | 0.100 | [6] |
| ABL | 0.100 to 0.200 | [8] |
| ABL | 0.100 to 0.300 | [5] |
| ABL | 0.110 | [10] |
| ABL | 0.156 ± 0.019 | [11] |

| | | |
|---|---|---|
| ABL | 0.190 | [12] |
| ABL | 0.267 | [13] |
| KIT | 0.052 | [10] |
| KIT | 0.099 | [14] |
| KIT | 0.100 | [3] |
| KIT | 0.100 | [5] |
| KIT | 0.100 | [15] |
| KIT | 0.124 | [16] |
| KIT | 0.137 | [12] |
| KIT | 0.300 | [9] |
| KIT | 0.400 | [7] |
| KIT | 0.400 | [6] |
| KIT | 0.410 | [4] |
| KIT | 0.410 | [8] |
| LCK | 0.320 | [10] |
| LCK | 0.600 to 0.800 | [17] |
| LCK | 0.900 | [9] |
| LCK | 1.000 | [6] |
| LCK | 2.600 | [18] |
| LCK | 9.000 | [4] |
| LCK | 9.000 | [8] |
| LCK | 9.000 | [5] |
| CHK1 | 1.000 | [19] |
| p38α | 10.00 to 72.44 | [21] |
| p38α | 13.70 | [10] |
| p38α | 70.00 | [6] |

| | | |
|---|---|---|
| SRC | 10.00 | [11] |
| SRC | 10.00 | [5] |
| SRC | 10.00 | [8] |
| SRC | 10.00 | [9] |
| SRC | 24.37 | [2] |
| SRC | 50.00 | [18] |
| SRC | 100.0 | [6] |
| SRC | 100.0 | [5] |
| SRC | 100.0 | [7] |
| SRC | 100.0 | [4] |

Table S3. Enzymatic affinity assays of imatinib across multiple kinases

| | $K_D$ (uM) | Reference Paper |
|---|---|---|
| ABL | 0.001 | [20] |
| ABL | 0.001 | [21] |
| ABL | 0.001 | [22] |
| ABL | 0.001 | [22] |
| ABL | 0.002 | [23] |
| ABL | 0.002 | [23] |
| ABL | 0.002 | [24] |
| ABL | 0.004 | [25] |
| ABL | 0.008 | [22] |
| ABL | 0.010 | [26] |
| ABL | 0.011 | [24] |

| | | |
|---|---|---|
| ABL | 0.012 | [27] |
| ABL | 0.012 | [6] |
| ABL | 0.014 | [28] |
| ABL | 0.024 | [23] |
| ABL | 0.034 | [29] |
| ABL | 0.080 | [7] |
| ABL | 0.1 | [30] |
| ABL | 0.106 | [31] |
| ABL | 0.282 | [25] |
| CHK1 | >3.0 | [21] |
| CHK1 | 10 | [30] |
| CHK1 | >10 | [27] |
| CHK1 | >10 | [24] |
| CHK1 | >30 | [31] |
| KIT | 0.003 | [22] |
| KIT | 0.007 | [22] |
| KIT | 0.013 | [32] |
| KIT | 0.013 | [20] |
| KIT | 0.013 | [21] |
| KIT | 0.013 | [24] |
| KIT | 0.014 | [27] |
| KIT | 0.110 | [29] |
| KIT | <0.1 | [30] |

| | | |
|---|---|---|
| LCK | 0.020 | [32] |
| LCK | 0.040 | [27] |
| LCK | 0.040 | [33] |
| LCK | 0.040 | [21] |
| LCK | 0.062 | [24] |
| LCK | 0.062 | [34] |
| LCK | <0.1 | [30] |
| p38α | 4.100 | [29] |
| p38α | 10 | [30] |
| p38α | >10 | [33] |
| p38α | >10 | [35] |
| p38α | >30 | [31] |
| p38α | 34.000 | [6] |
| SRC | >10 | [33] |
| SRC | >10 | [24] |
| SRC | >10 | [7] |
| SRC | >30 | [31] |
| SRC | 42.000 | [36] |

Table S4. Upper and Lower Bound Calculations from $E_{tot}$ enthalpy and protein binding pocket dihedral entropy for all protein systems

| Kinase | $\Delta H_{upper}$ | $\Delta H_{lower}$ | $T\Delta S_{upper}$ | $T\Delta S_{lower}$ | $\Delta G_{upper}$ | $\Delta G_{lower}$ |
|---|---|---|---|---|---|---|
| ABL | -53.38 | -82.19 | -6.68 | -10.56 | -42.83 | -75.51 |
| KIT | -56.90 | -82.71 | -4.78 | -6.63 | -50.27 | -77.94 |
| LCK | -28.17 | -47.86 | -1.35 | -11.21 | -16.96 | -46.50 |
| CHK1 | -13.82 | -40.66 | -0.53 | -5.68 | -8.15 | -40.14 |
| p38α | -15.97 | -30.32 | -1.83 | -5.54 | -10.43 | -28.49 |
| SRC | -21.47 | -37.71 | -2.23 | -11.03 | -10.44 | -35.48 |

To obtain the specific values shown in the cells, please refer to the supplemental data code for the changes in enthalpy, entropy, and Gibbs energy calculations. For the method on how to calculate $\Delta H_{upper}$, $\Delta H_{lower}$, $\Delta S_{upper}$, $\Delta S_{lower}$, $\Delta G_{upper}$, and $\Delta G_{lower}$, please refer to the "Enthalpy and Dihedral Entropy Calculations" section of the main manuscript. To calculate for the upper-bound $\Delta G_{upper}$, take the difference between $\Delta H_{upper}$ and $T\Delta S_{lower}$. To calculate for the upper-bound $\Delta G_{lower}$, take the difference between $\Delta H_{lower}$ and $T\Delta S_{upper}$. This sets the possible boundaries for a fair comparison with the $K_D$ and $IC_{50}$ ranges compiled in table S2 and table S3.

Figure S1. Bound-Complex Kinase MM/PBSA analysis reveals converging interaction energies of selected kinases. The first 50 ns are ignored to account for equilibration. The first, second, and third repeats are shown in blue, red, and yellow, respectively.

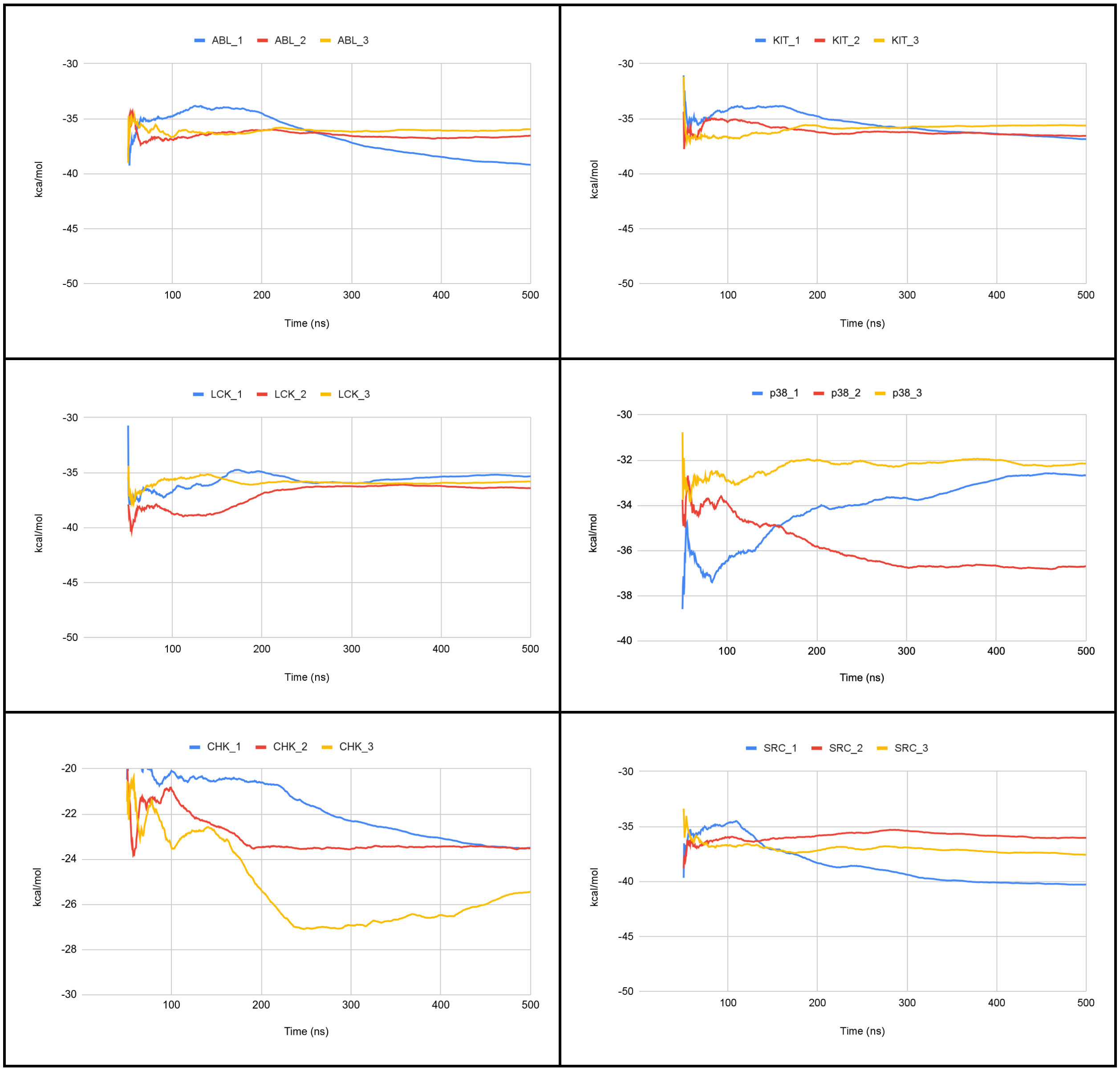

Figure S2. $E_{tot}$ enthalpy convergences calculated with R. The specific convergence intervals are system-dependent and included in the supplemental data folder.

ABL bound seeds 1, 2, and 3

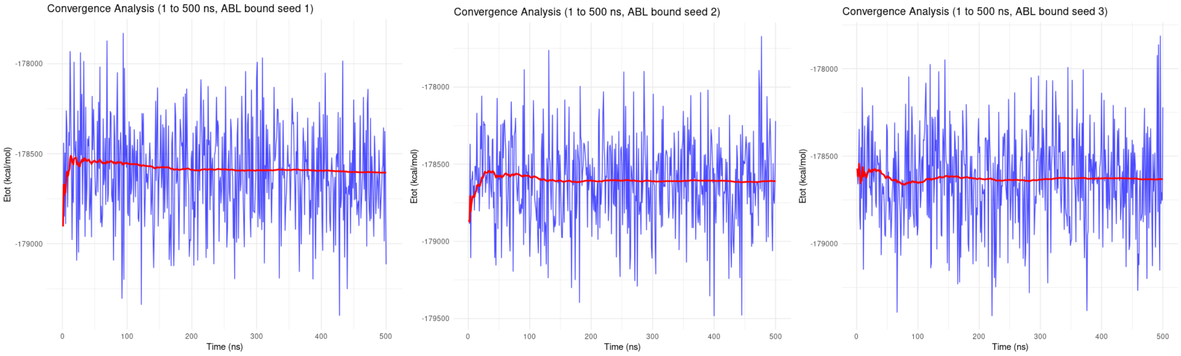


ABL free seeds 1, 2, and 3

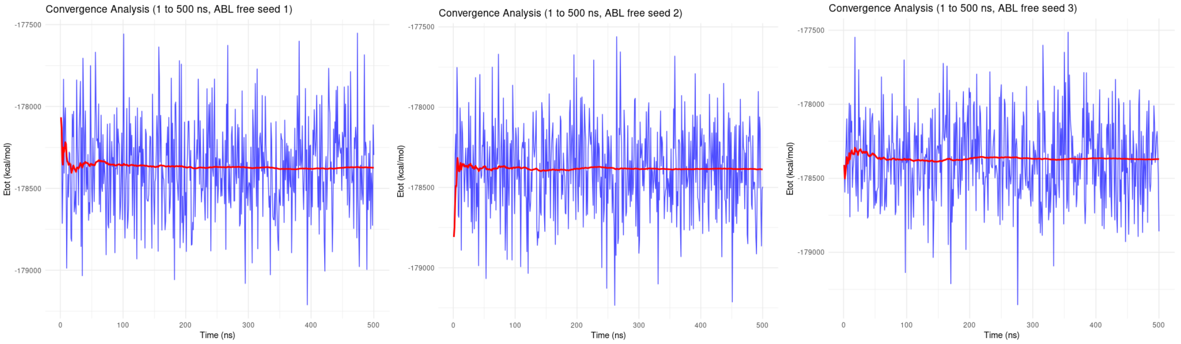


KIT bound seeds 1, 2, and 3

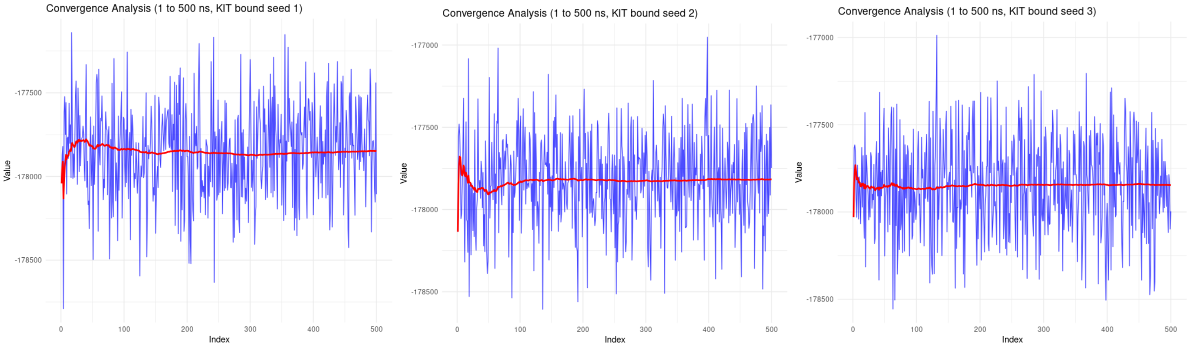


KIT free seeds 1, 2, and 3

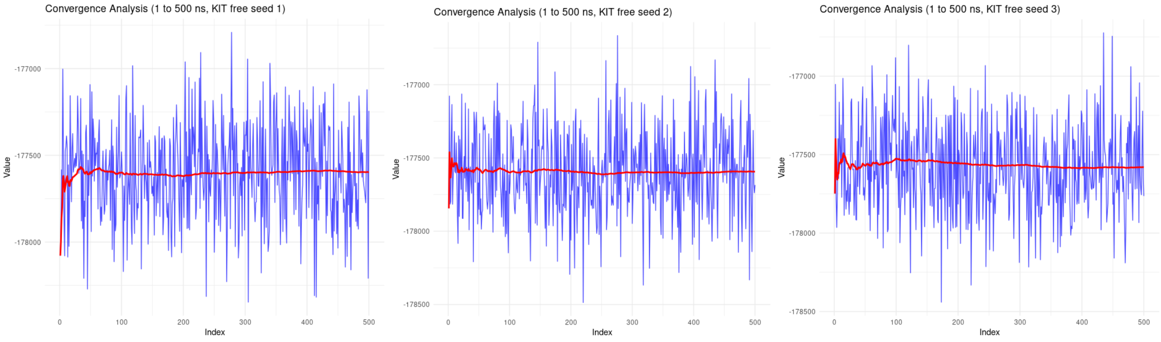


LCK bound seeds 1, 2, and 3

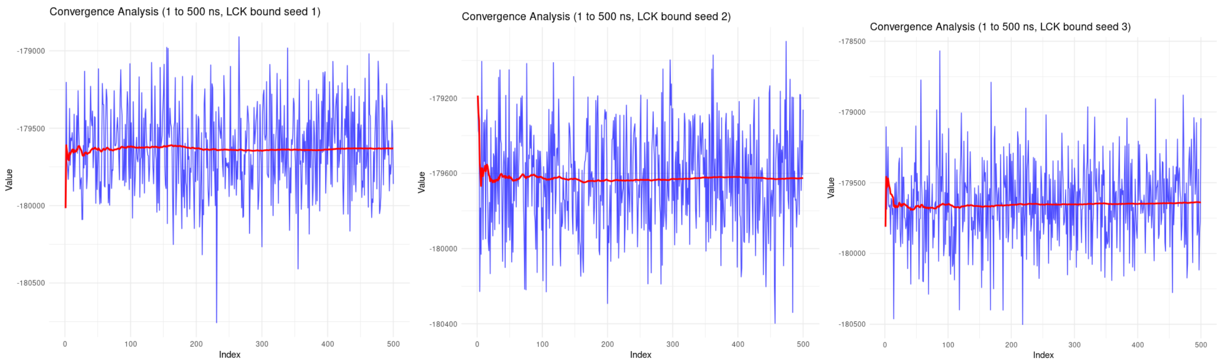


LCK free seeds 1, 2, and 3

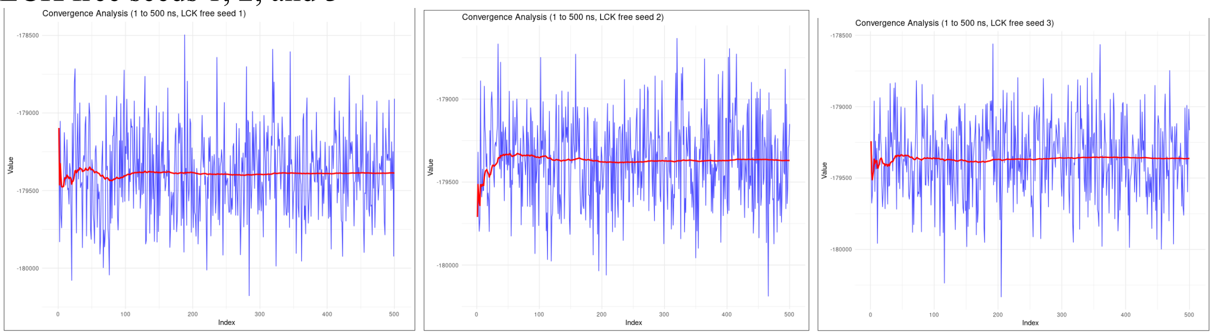


CHK1 bound seeds 1, 2, and 3

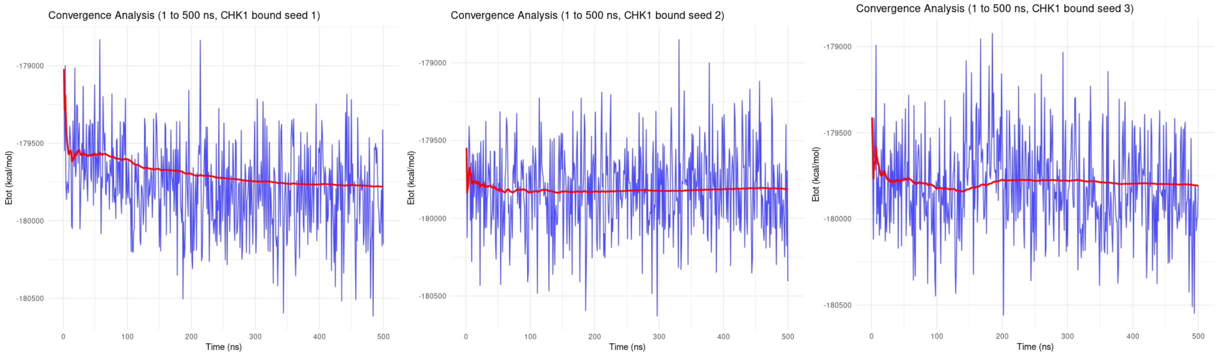


CHK1 free seeds 1, 2, and 3

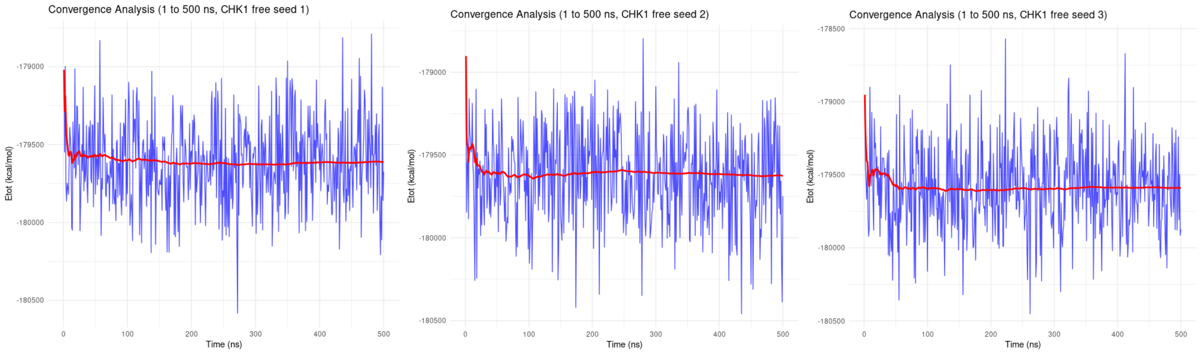

### p38α bound seeds 1, 2, and 3

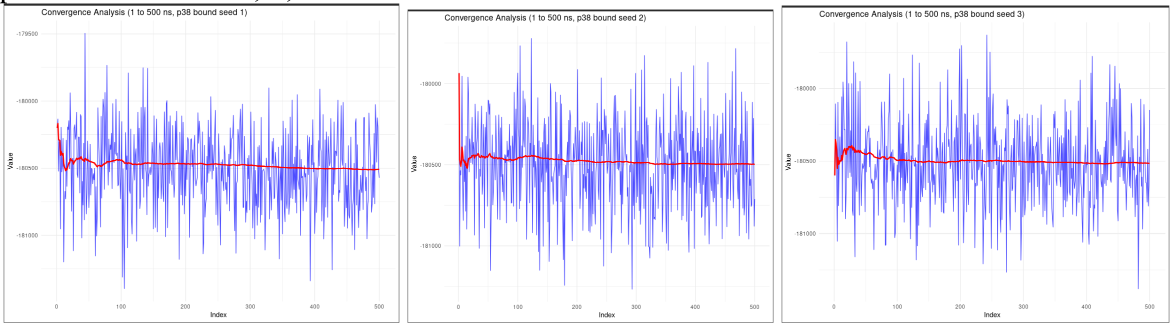


### p38α free seeds 1, 2, and 3

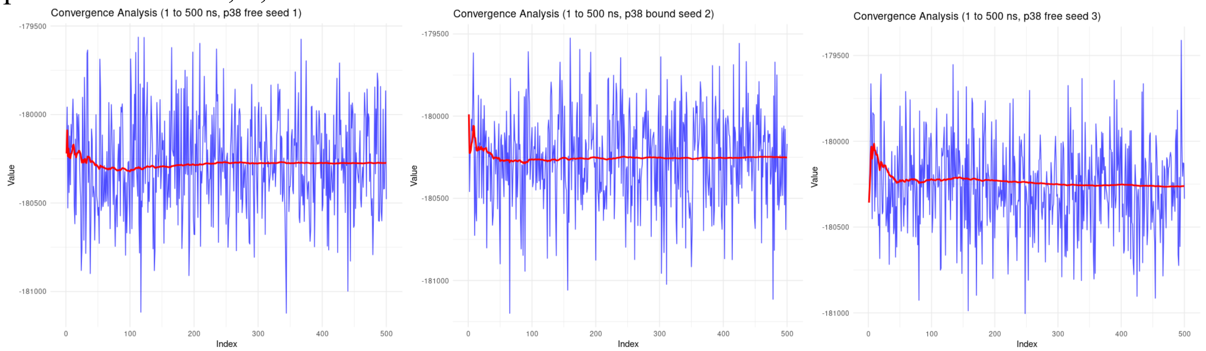


### SRC bound seeds 1, 2 and 3

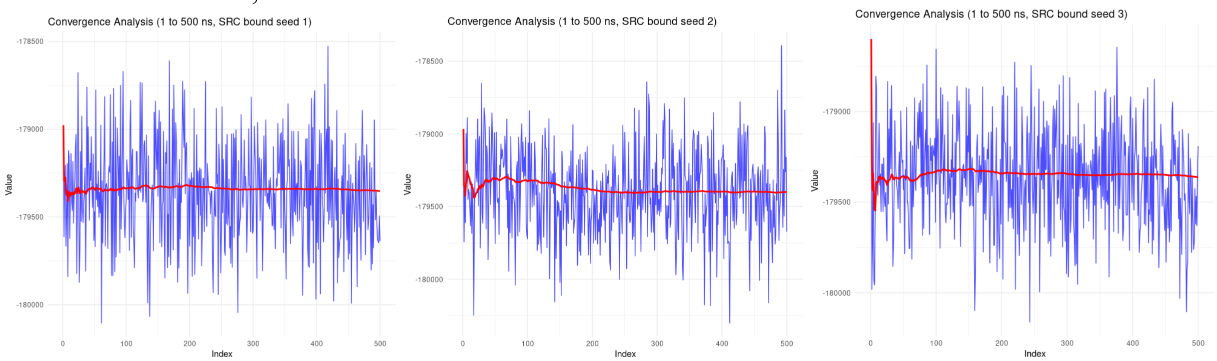


### SRC free seeds 1, 2, and 3

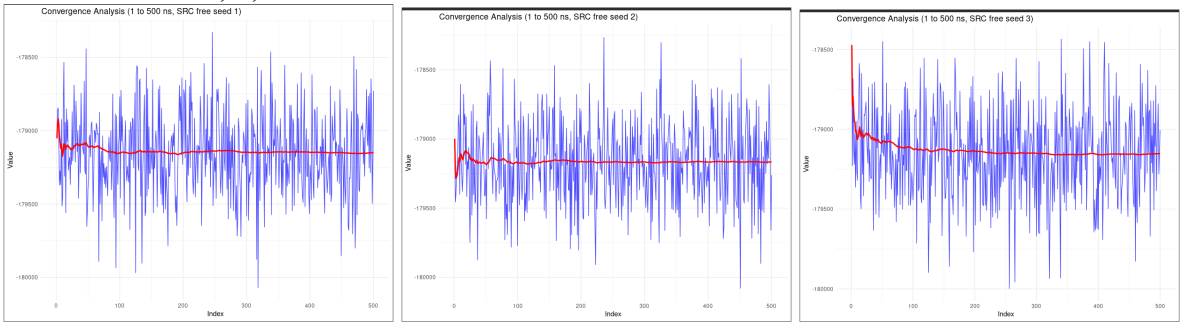

### Imatinib free seeds 1, 2, and 3

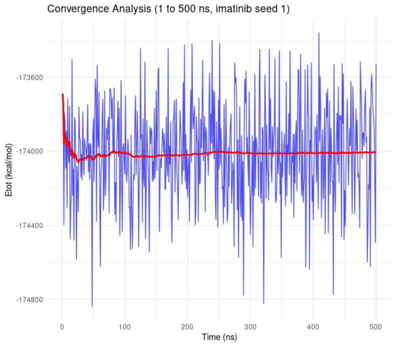


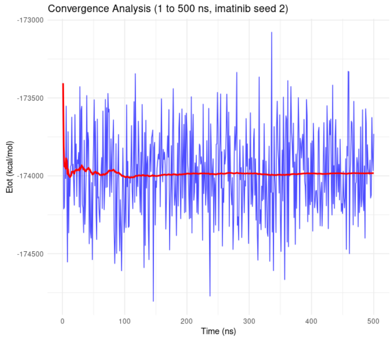


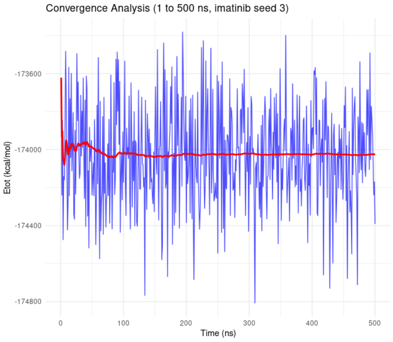


### Waterbox seeds 1, 2, and 3

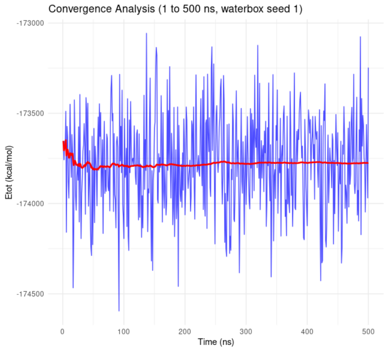


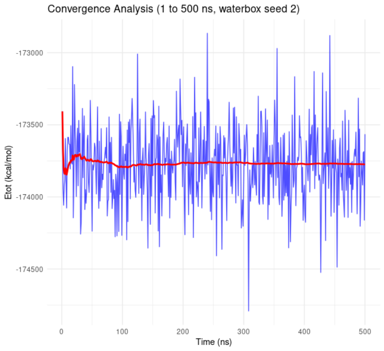


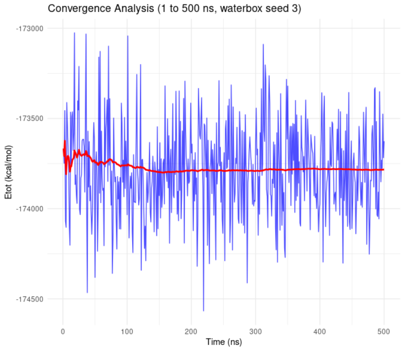

Figure S3: Root-mean-square-deviation (RMSD) stability quantification. The free protein plots are shown on the left, while the bound protein plots are shown to the right. The first, second, and third repeats are shown in blue, red, and yellow, respectively.

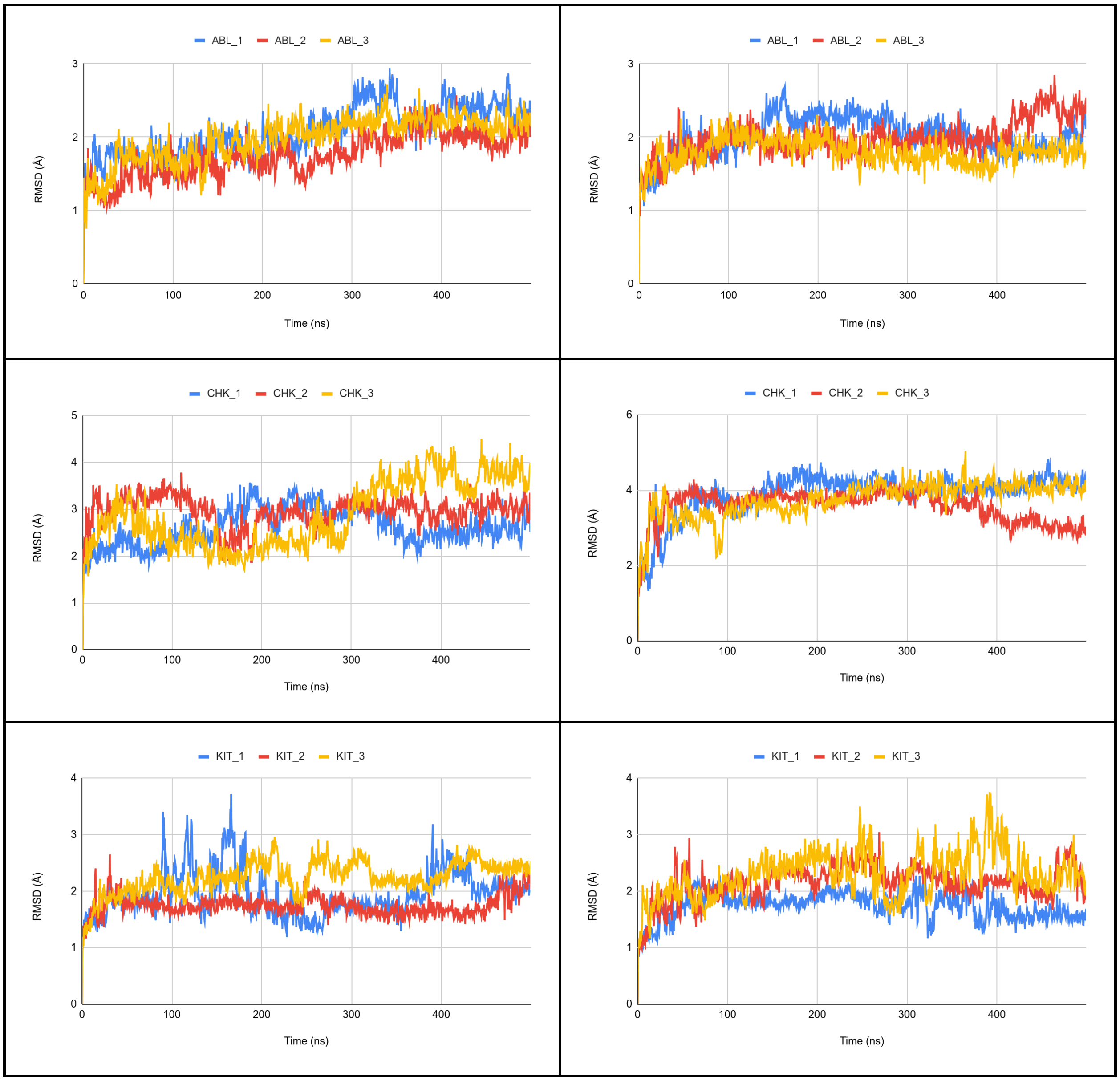

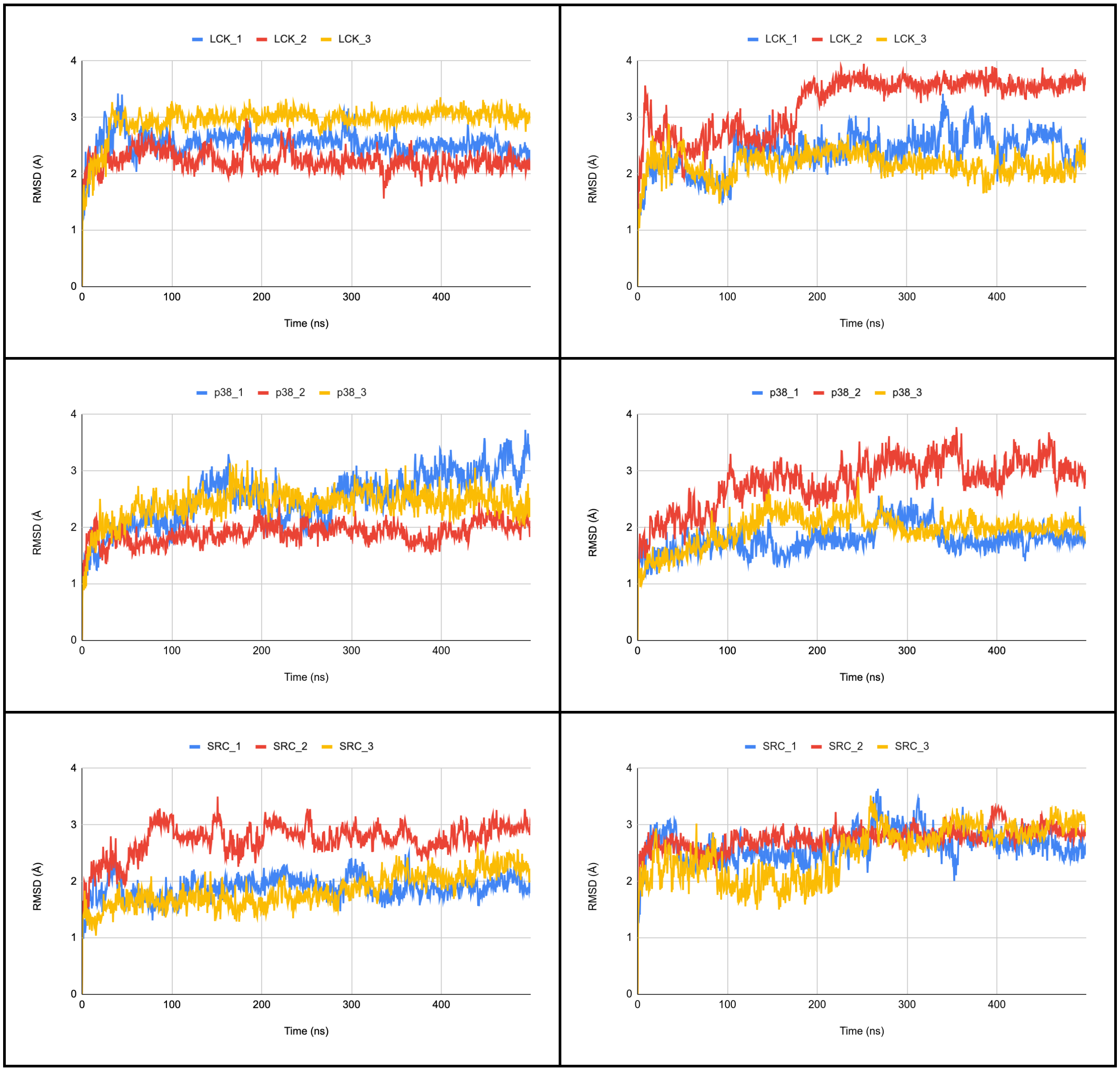
LCK_1 LCK_2 LCK_3
RMSD (A)
Time (ns)
LCK_1 LCK_2 LCK_3
RMSD (Å)
Time (ns)
p38_1 p38_2 p38_3
RMSD (Å
Time (ns)
p38_1 p38_2 p38_3
RMSD (Å)
Time (ns)
SRC_1 SRC_2 SRC_3
RMSD (A)
Time (ns)
SRC_1 SRC_2 SRC_3
RMSD (Å)
Time (ns)

Figure S4. Root-mean-square-fluctuation (RMSF) stability quantification. The free protein plots are shown to the left, while the bound protein plots are shown to the right. The first, second, and third repeats are shown in blue, red, and yellow, respectively.

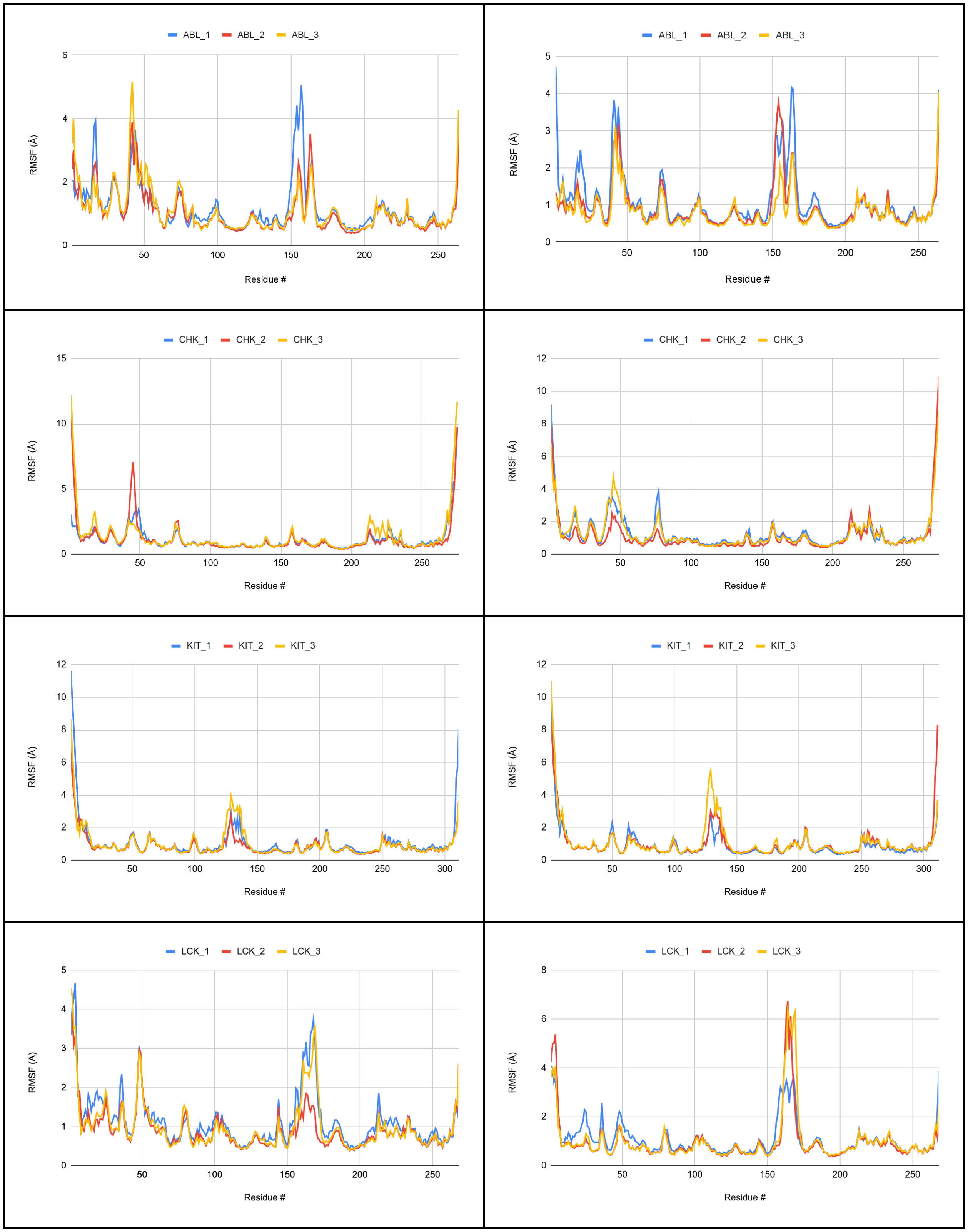

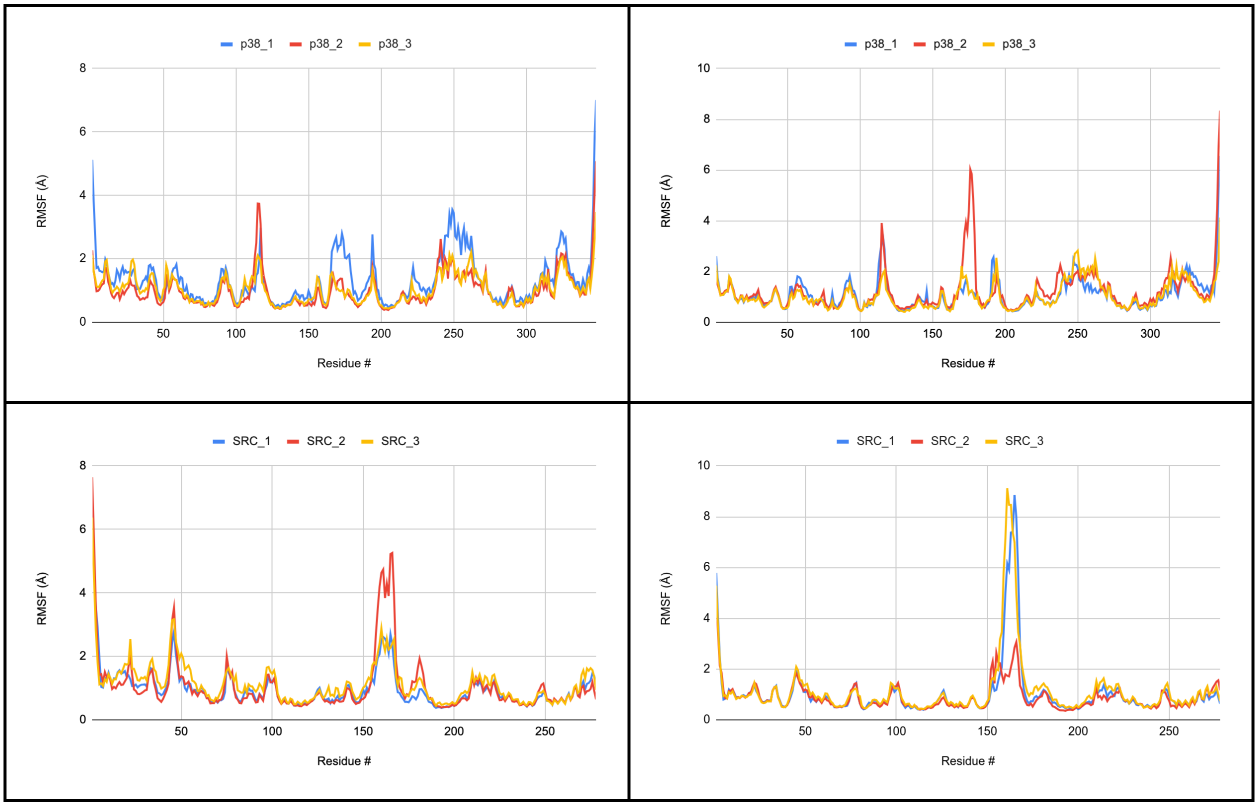

p38_1 p38_2 p38_3
RMSF (Å)
Residue #
SRC_1 SRC_2 SRC_3
RMSF (Å)
Residue #

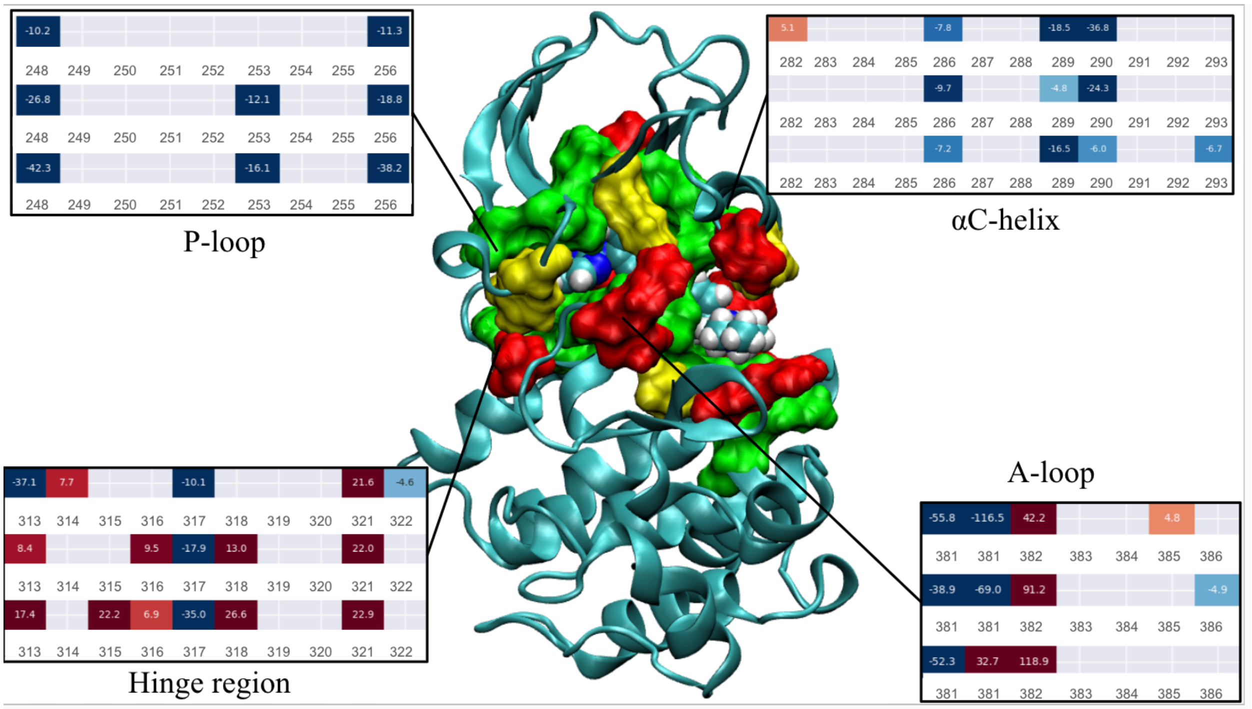


Figure S5. Force distribution analysis and the corresponding 3D region for the ABL kinase. In the boxes, blue indicates attractive interactions, while red indicates repulsive interactions. Residues exhibiting ~ ±5 pN force on average are shown in surface. Green, yellow, and red exhibit, on average, more than ±5 pN of force in three, two, and one MD runs, respectively.

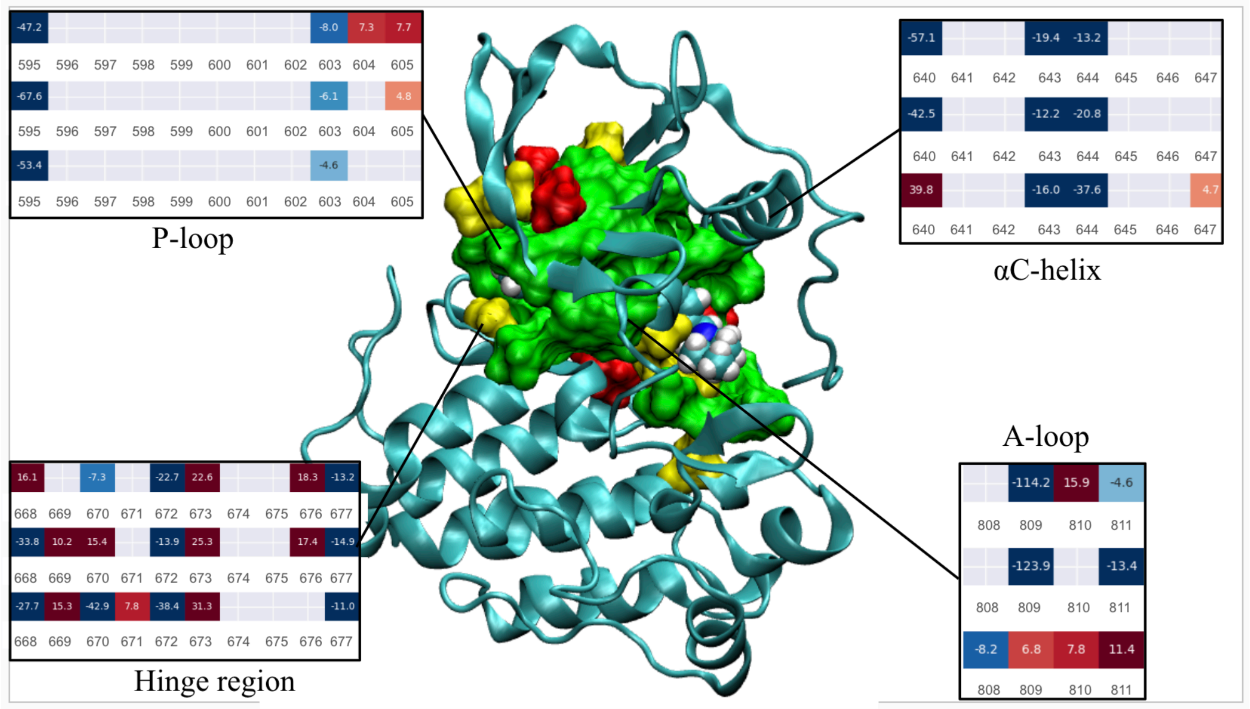


Figure S6. Force distribution analysis and the corresponding 3D region for the KIT kinase. In the boxes, blue indicates attractive interactions, while red indicates repulsive interactions. Residues exhibiting ~ ±5 pN force on average are shown in surface. Green, yellow, and red exhibit, on average, more than ±5 pN of force in three, two, and one MD runs, respectively.

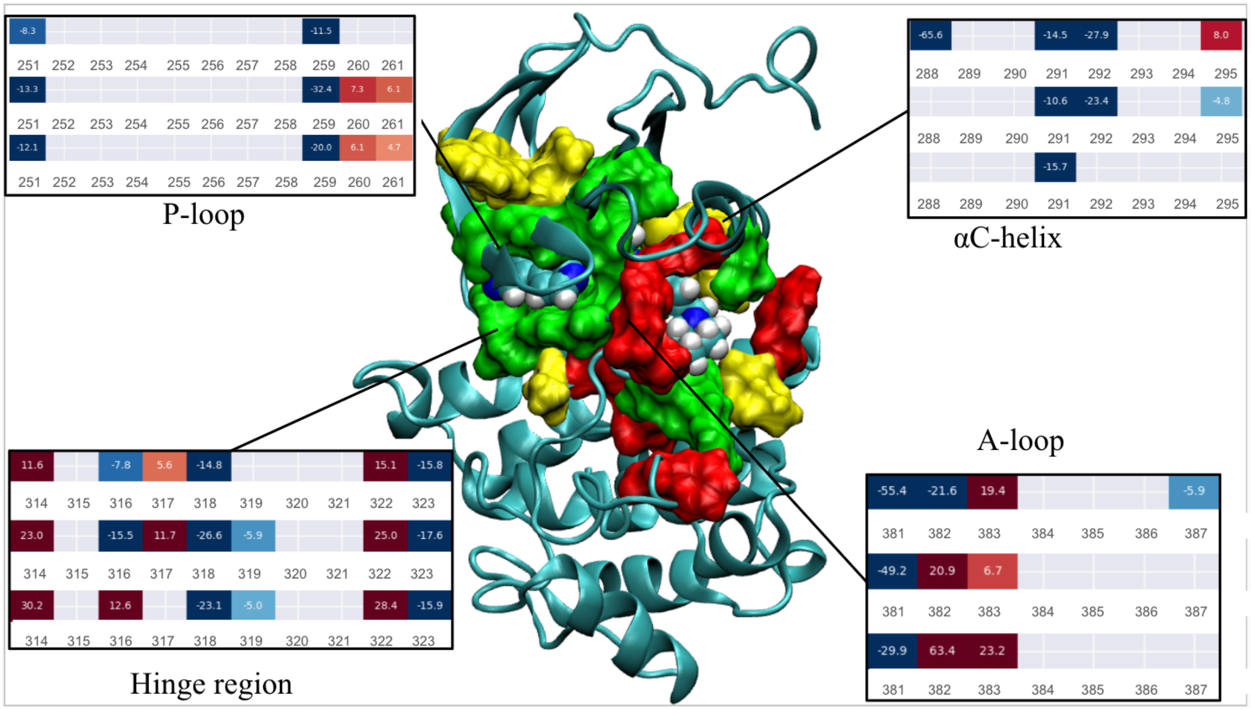


Figure S7. Force distribution analysis and the corresponding 3D region for the Lck kinase. In the boxes, blue indicates attractive interactions, while red indicates repulsive interactions. Residues exhibiting ~ ±5 pN force on average are shown in surface. Green, yellow, and red exhibit, on average, more than ±5 pN of force in three, two, and one MD runs, respectively.

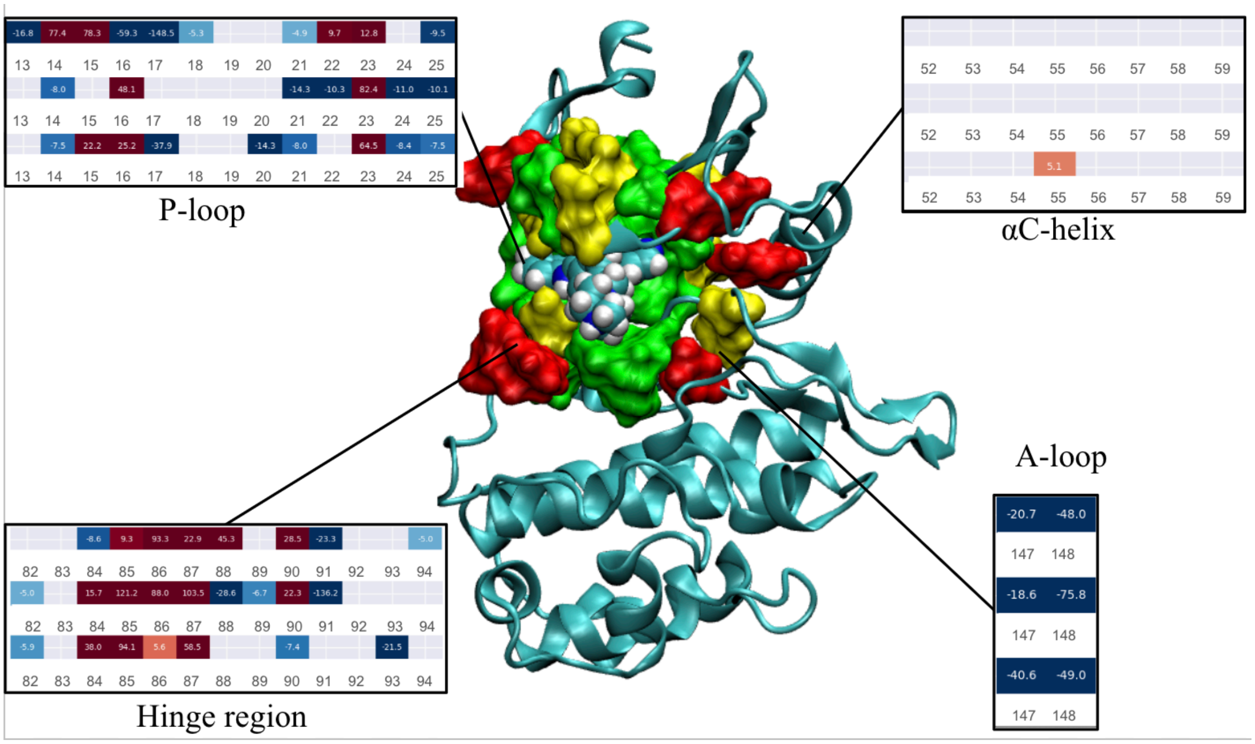


Figure S8. Force distribution analysis and the corresponding 3D region for the Chk1 kinase. In the boxes, blue indicates attractive interactions, while red indicates repulsive interactions. Residues exhibiting ~ ±5 pN force on average are shown in surface. Green, yellow, and red exhibit, on average, more than ±5 pN of force in three, two, and one MD runs, respectively.

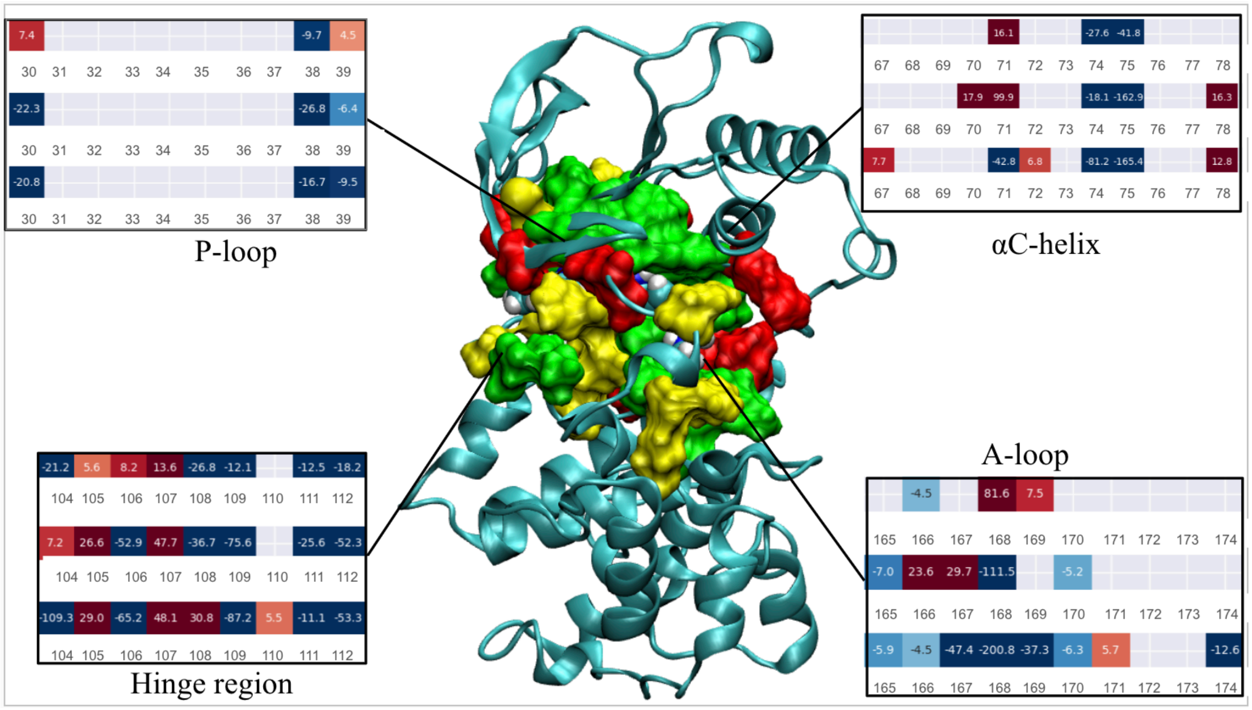


Figure S9. Force distribution analysis and the corresponding 3D region for the p38α kinase. In the boxes, blue indicates attractive interactions, while red indicates repulsive interactions. Residues exhibiting ~ ±5 pN force on average are shown in surface. Green, yellow, and red exhibit, on average, more than ±5 pN of force in three, two, and one MD runs, respectively.

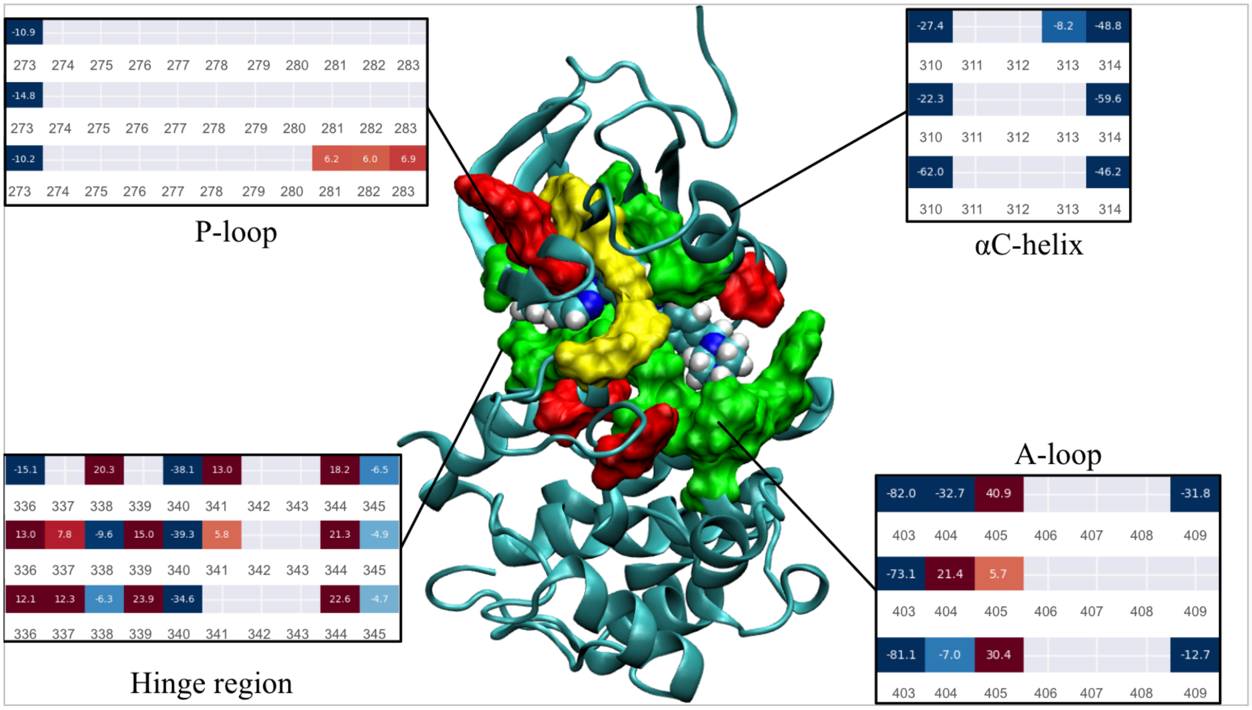


Figure S10. Force distribution analysis and the corresponding 3D region for the Src kinase. In the boxes, blue indicates attractive interactions, while red indicates repulsive interactions. Residues exhibiting ~ ±5 pN force on average are shown in surface. Green, yellow, and red exhibit, on average, more than ±5 pN of force in three, two, and one MD runs, respectively.

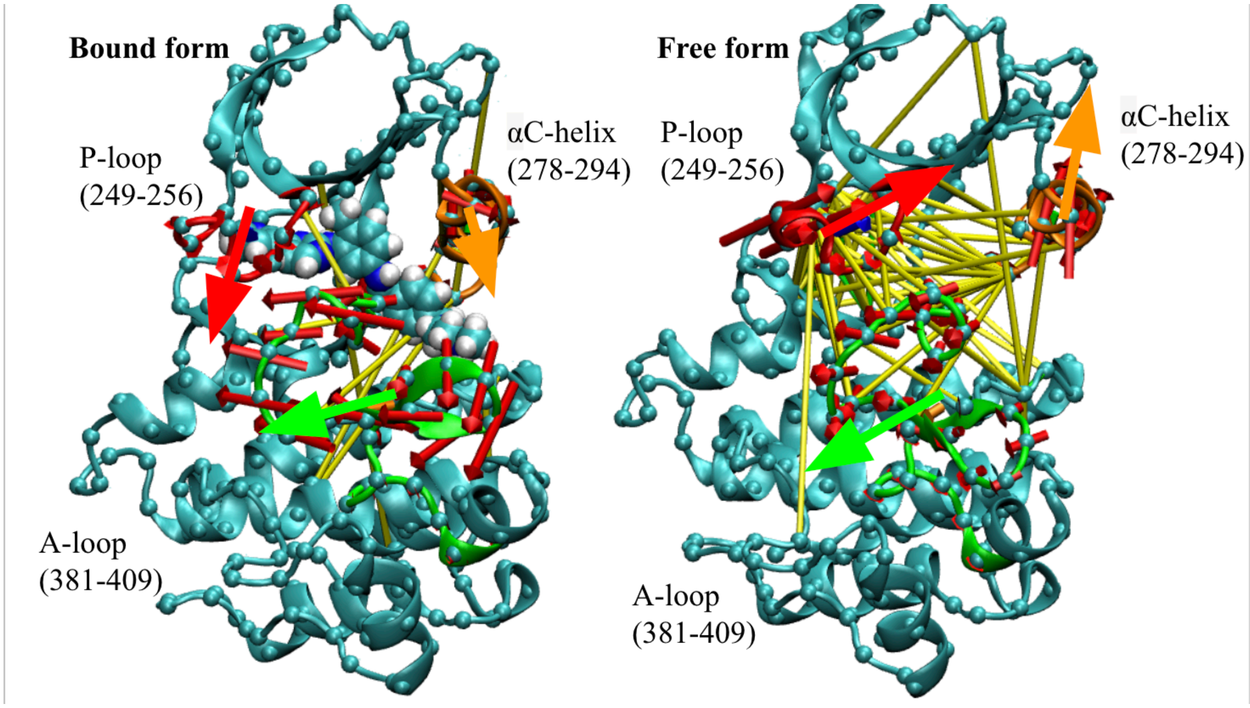


Figure S11. Principal component analysis and the sidechain network correlation visualization for the Abl kinase. The yellow cylinders correspond with an (anti-)correlation cutoff of ±0.3 in two experimental repeats. The P-loop motions are shown in red, the αC-helix in orange, and the A-loop in green. The large red, orange, and green arrows show the average migration pattern for the secondary motif based on the eigenvectors for the first PC calculated from BKiT. The snapshots are captured at the end of a 500 ns trajectory.

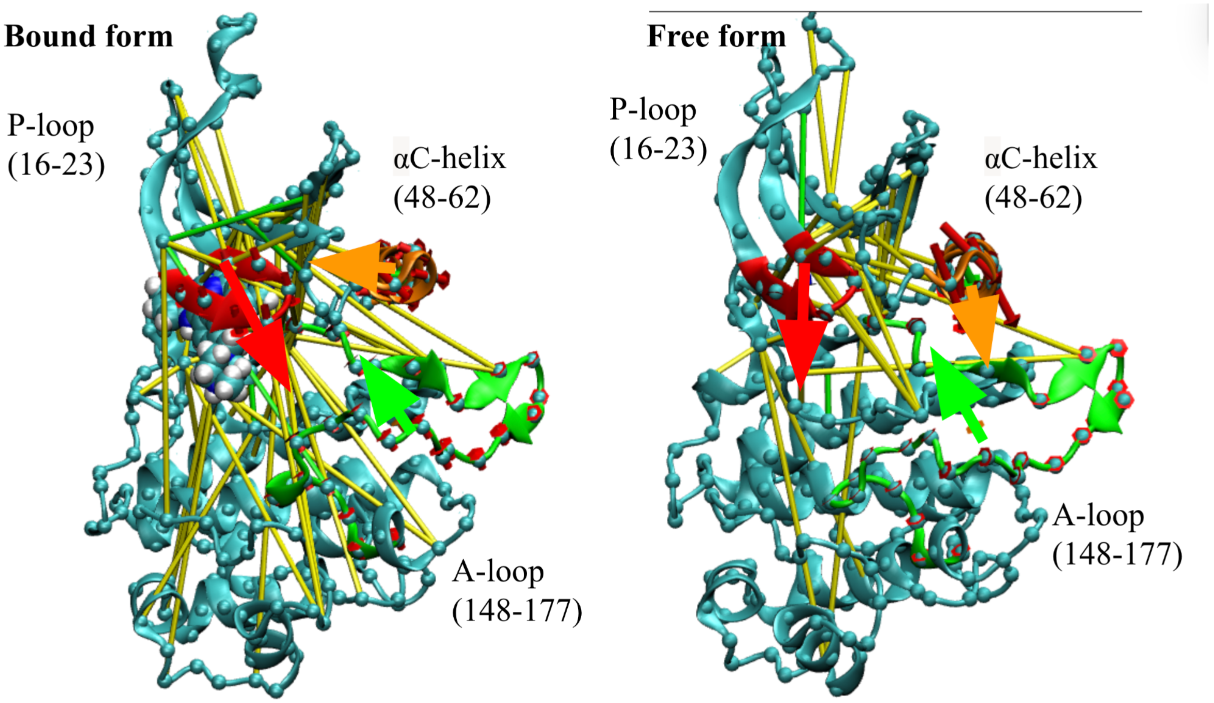


Figure S12. Principal component analysis and the sidechain network correlation visualization for the Chk1 kinase. Red is the P-loop, orange is the αC-helix, and green is the A-loop. The yellow cylinders correspond with an (anti-)correlation cutoff of ±0.3 in two experimental repeats. The P-loop motions are shown in red, the αC-helix in orange, and the A-loop in green. The large red, orange, and green arrows show the average migration pattern for the secondary motif based on the eigenvectors for the first PC calculated from BKiT. The snapshots are captured at the end of a 500 ns trajectory.

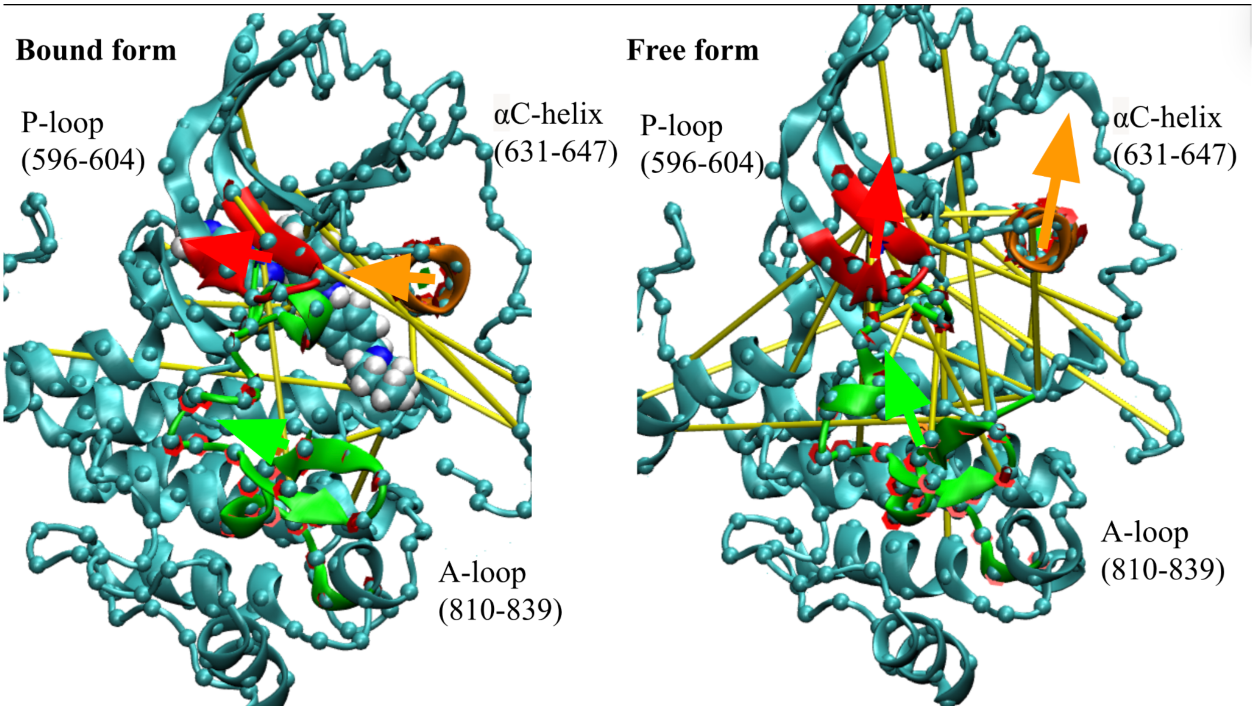


Figure S13. Principal component analysis and the sidechain network correlation visualization for the Kit kinase. Red is the P-loop, orange is the αC-helix, and green is the A-loop. The yellow cylinders correspond with an (anti-)correlation cutoff of ±0.3 in two experimental repeats. The P-loop motions are shown in red, the αC-helix in orange, and the A-loop in green. The large red, orange, and green arrows show the average migration pattern for the secondary motif based on the eigenvectors for the first PC calculated from BKiT. The snapshots are captured at the end of a 500 ns trajectory.

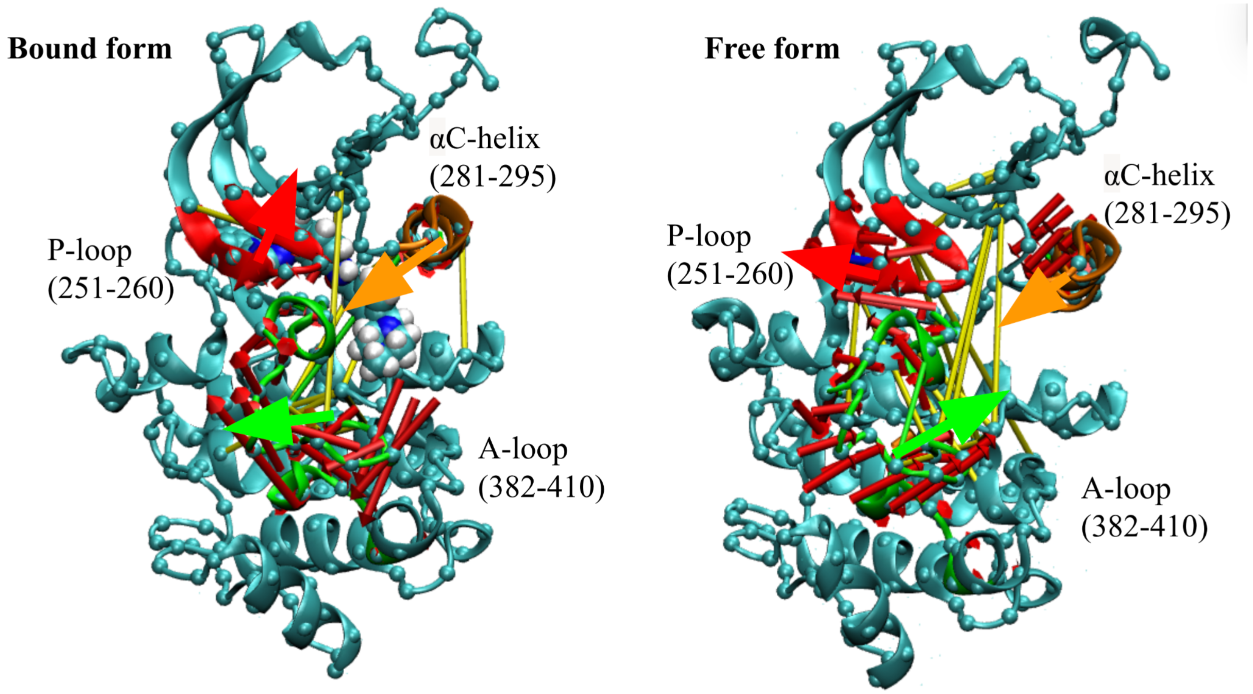


Figure S14. Principal component analysis and the sidechain network correlation visualization for the Lck kinase. Red is the P-loop, orange is the αC-helix, and green is the A-loop. The yellow cylinders correspond with an (anti-)correlation cutoff of ±0.3 in two experimental repeats. The P-loop motions are shown in red, the αC-helix in orange, and the A-loop in green. The large red, orange, and green arrows show the average migration pattern for the secondary motif based on the eigenvectors for the first PC calculated from BKiT. The snapshots are captured at the end of a 500 ns trajectory.

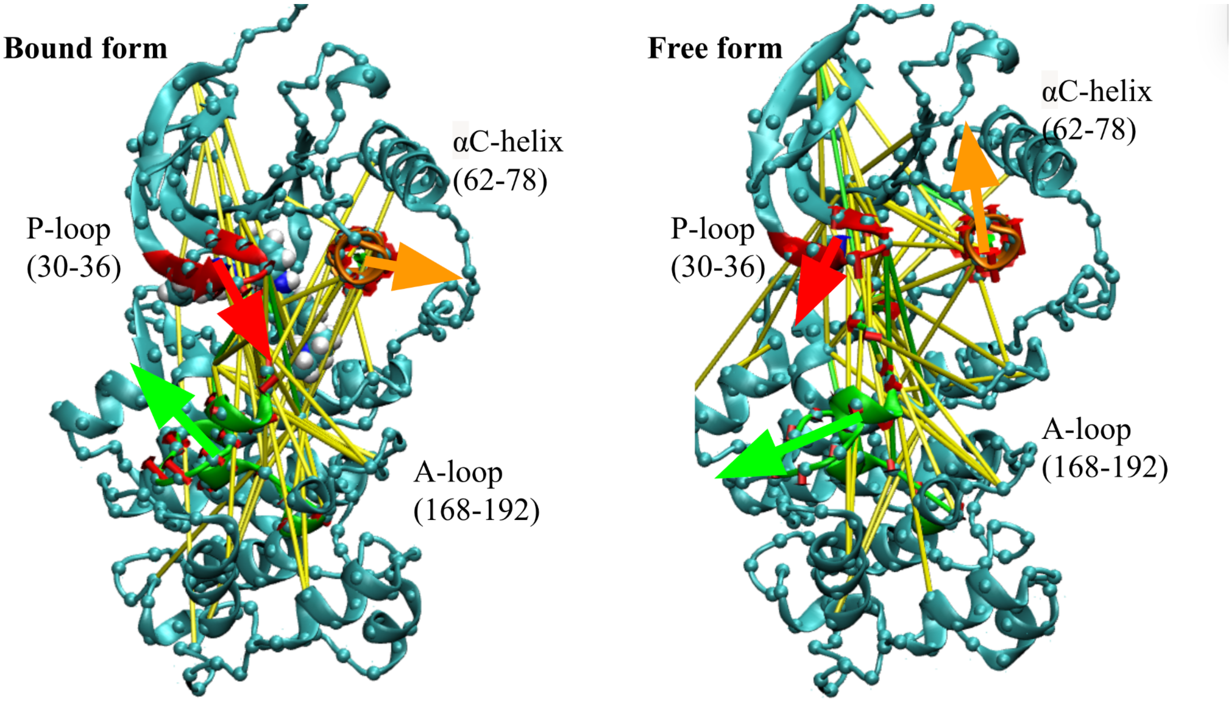


Figure S15. Principal component analysis and the sidechain network correlation visualization for the p38α kinase. Red is the P-loop, orange is the αC-helix, and green is the A-loop. The yellow cylinders correspond with an (anti-)correlation cutoff of ±0.3 in two experimental repeats. The P-loop motions are shown in red, the αC-helix in orange, and the A-loop in green. The large red, orange, and green arrows show the average migration pattern for the secondary motif based on the eigenvectors for the first PC calculated from BKiT. The snapshots are captured at the end of a 500 ns trajectory.

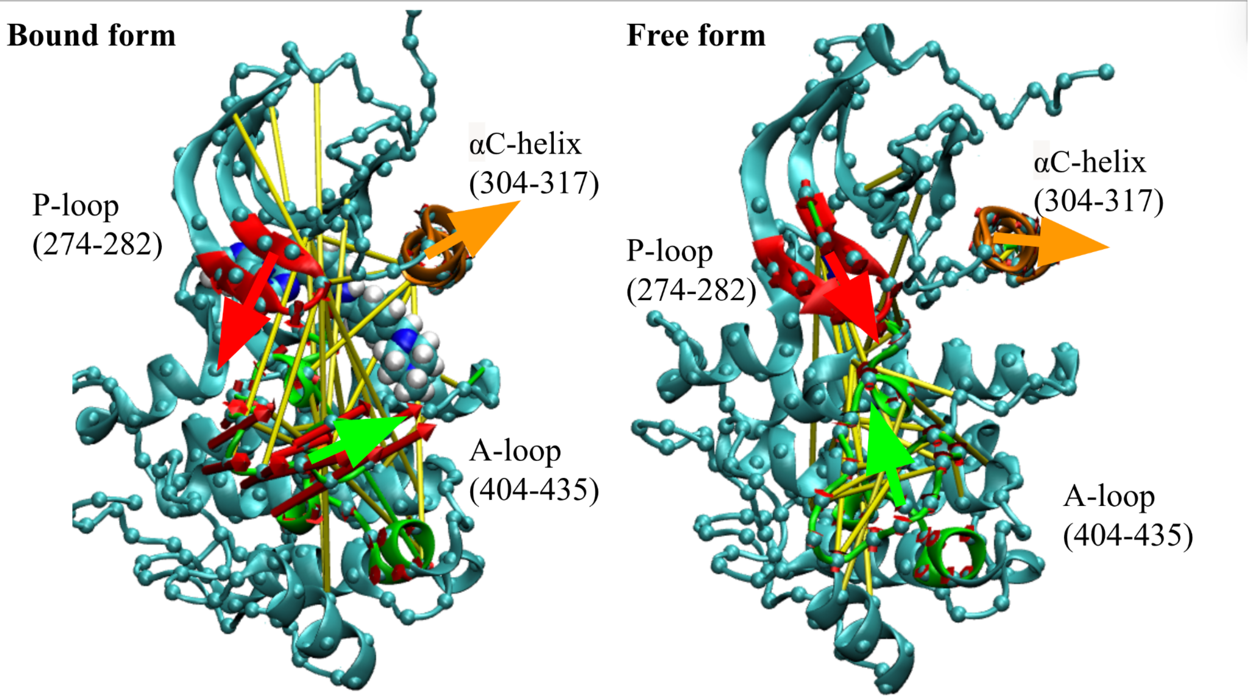


Figure S16. Principal component analysis and the sidechain network correlation visualization for the Src kinase. Red is the P-loop, orange is the αC-helix, and green is the A-loop. The yellow cylinders correspond with an (anti-)correlation cutoff of ±0.3 in two experimental repeats. The P-loop motions are shown in red, the αC-helix in orange, and the A-loop in green. The large red, orange, and green arrows show the average migration pattern for the secondary motif based on the eigenvectors for the first PC calculated from BKiT. The snapshots are captured at the end of a 500 ns trajectory.
